# Contribution of Apical and Basal Dendrites of L2/3 Pyramidal Neurons to Orientation Encoding in Mouse V1

**DOI:** 10.1101/566588

**Authors:** Jiyoung Park, Athanasia Papoutsi, Ryan T. Ash, Miguel A. Marin, Panayiota Poirazi, Stelios M. Smirnakis

## Abstract

Pyramidal neurons integrate synaptic inputs from basal and apical dendrites to generate stimulus-specific responses. It has been proposed that feed-forward inputs to basal dendrites drive a neuron’s stimulus preference, while feedback inputs to apical dendrites sharpen selectivity. However, how a neuron’s dendritic domains relate to its functional selectivity has not been demonstrated experimentally. We performed 2-photon dendritic micro-dissection on layer-2/3 pyramidal neurons in mouse primary visual cortex. We found that removing the apical dendritic tuft did not alter orientation-tuning. Furthermore, orientation-tuning curves were remarkably robust to the removal of basal dendrites: ablation of 2-3 basal dendrites was needed to cause a small shift in orientation preference, without significantly altering tuning width. Computational modeling corroborated our results and put limits on how orientation preferences among basal dendrites differ in order to reproduce the post-ablation data. In conclusion, neuronal orientation-tuning appears remarkably robust to loss of dendritic input.

---

Neocortical pyramidal neurons ramify several basal dendritic arbors laterally and one apical dendritic arbor superficially to receive and integrate synaptic inputs^1^. Each dendritic-tree of a mouse layer 2/3 (L2/3) pyramidal neuron arborizes in a different cortical map subregion (e.g. retinotopic region) and/or cortical layer, thereby sampling largely non-overlapping axonal inputs coming in from different brain areas^2^. Inputs are functionally heterogeneous across dendrites and even within individual dendritic branches^3,4^. Mouse primary visual cortex (V1) L2/3 pyramidal neurons generate action potentials in response to a narrow range of orientations^5^ despite receiving highly heterogeneous input and poorly tuned subthreshold responses^4^, making them ideal for studying the relationship between dendritic input and functional selectivity. Indeed, recent evidence suggests that apical tuft dendritic spikes serve to narrow the orientation tuning function, increasing orientation selectivity of area V1 L2/3 pyramidal neurons^6^. L2/3 pyramidal neurons are also a good model system for studying the relative roles of apical and basal dendrites: while basal dendrites primarily receive feedforward input from L4 and nearby L2/3 neurons^7,8^, apical dendrites receive cortico-cortical feedback that may refine orientation selectivity^9,10^ as well as orientation-tuned thalamo-cortical input from layer 1^4,11–13^.

It is difficult to firmly establish a causal relationship between computations that occur in dendritic branches and properties at the soma. Calcium imaging does not always have the temporal resolution to disambiguate signals arising in different dendritic processes or the soma, while *in vivo* dendritic patch clamping, despite its power at dissecting specific hypotheses, typically disrupts normal dendro-somatic processing making it difficult to dissect dendritic influences on the soma. The existence of functional inputs and activity on dendrites does not explain the necessity of these inputs to the neuron’s final output. Here we employed *in vivo* two photon microdissection^14–17^ to systematically remove individual dendrites from layer 2/3 mouse V1 pyramidal neurons, allowing us to assess the causal relationship between inputs arriving in different dendritic arbors and the computation of orientation selectivity at the soma.

## RESULTS

### Dendrite Ablation *in vivo*

To assess the causal relationship between inputs arriving in different dendrites and orientation-tuning, we systematically removed individual dendrites from L2/3 mouse V1 pyramidal neurons using *in vivo* 2-photon micro-dissection^14,15,17^. We visualized both dendritic structure and functional activity of L2/3 pyramidal neurons by stereotactically co-injecting AAV-flex-GCaMP6s and diluted AAV-CaMKII-Cre (40,000 to 120,000X) into L2/3 of mouse V1^11^ (Fig. 1a, methods). Individual dendrites were then removed by performing 2-photon laser point scans on the targeted fluorescent dendritic segment, 10-30 μm away from the soma (Fig. 1b,c and Supp. Fig. 1)^14,15^. Dendritic segments expressing GCaMP6s, located 130-200 μm from the dura, could be typically ablated with one or two 200-400 ms point scans of 140-200 mW at 800 nm. Ablated dendrites distal to the lesion acquired a beads-on-the-string appearance and degraded, disappearing within 1-3 hours (Fig. 1b, Supp. Movie 1,2)^14,15^. Upon ablation, target neurons transiently became brightly fluorescent (Supp. Fig. 1d), presumably due to the influx of calcium through the instantaneous opening in the membrane, returning to pre-ablation fluorescence levels over the next 5-90 minutes. Neurons whose fluorescence did not return to its pre-ablation levels within 90 minutes were more likely to be visually undetectable the next day. Neurons whose apical dendritic arbors were ablated immediately adjacent to the soma were also more likely to disappear, so we restricted ablation to neurons with an extended primary apical trunk (≥ 20-30 μm from soma to the first apical bifurcation), targeting the point immediately prior to the bifurcation. Neurons that were still present 24 hours post-ablation (42% of cells: 11 out of 26 neurons from 19 mice survived after apical dendrite ablation, see Methods) exhibited intact morphology in their residual (non-ablated) dendritic segments and normal spontaneous and visual-evoked somatic calcium transients (Fig. 1b-d, Fig. 2a and Supp. Fig. 1).

**Figure 1.**
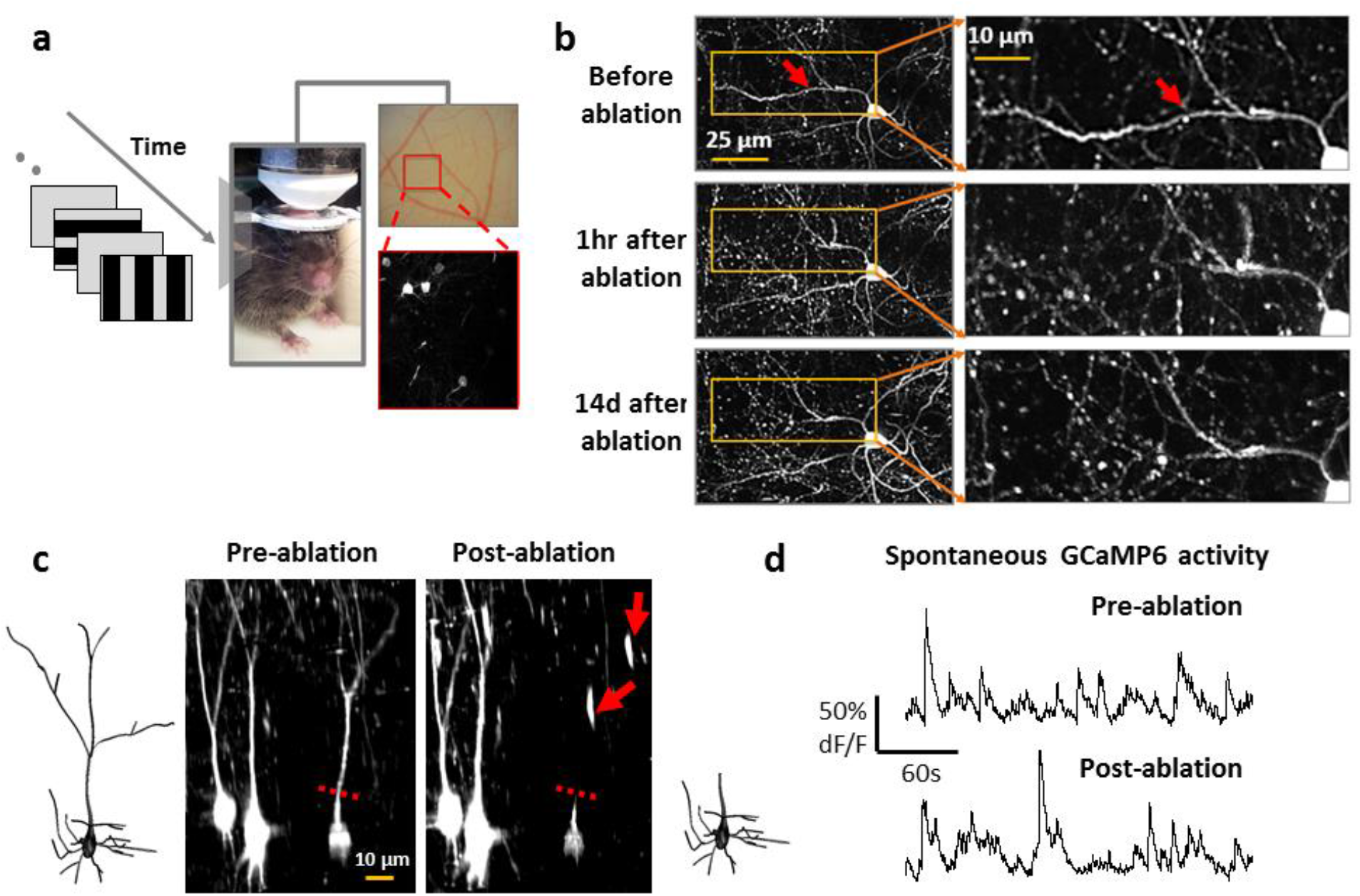
*In vivo* 2-photon laser dissection of apical dendrites in mouse primary visual cortex. **(a)** Experiment set-up. *Left:* visual stimulus. *<u>Middle</u>:* mouse under the objective. *<u>Right top</u>:* optical image of the surface of V1. *<u>Right bottom:</u>* two-photon image acquired from the inset showing sparse GCaMP6s-labeled neurons. (b) Dendritic arbor before (top), 1 hour (middle) and 14 days (bottom) after targeted laser-dissection of a single dendritic branch (red arrow: ablation point). Right panels zoom in. (c) *<u>Center:</u>* 3D renderings of the target neuron and its neighboring control neurons before (left) and after (right) apical dendrite ablation. Broken red line: the ablation point. Red arrows: GCaMP filled remnants of the severed apical dendrite. 3D reconstruction of target neuron before (far left) and after (far right) apical dendrite ablation. (d) Spontaneous GCaMP6s activity of the example neuron before and after apical tuft ablation.

**Figure 2.**
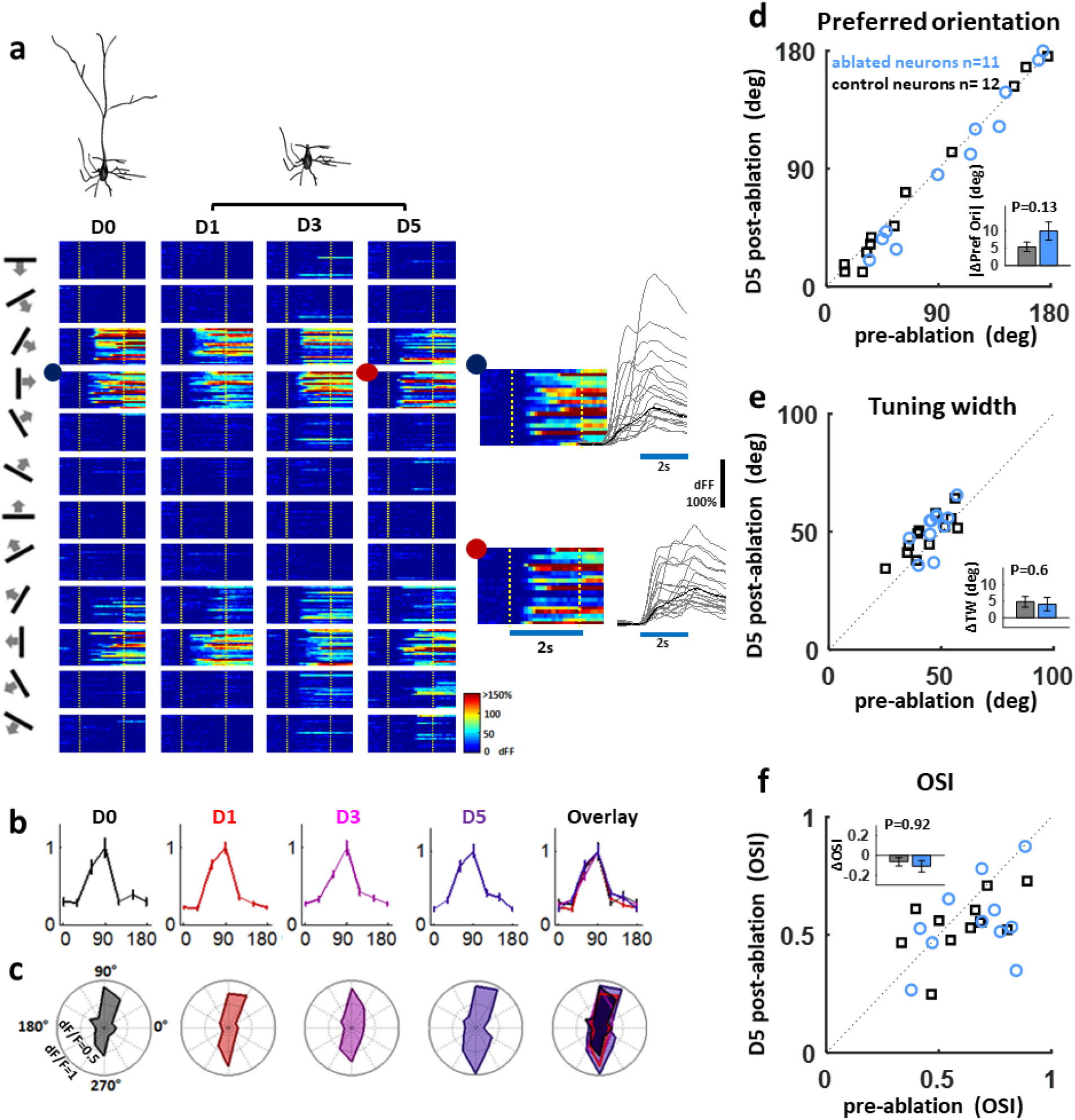
Orientation-tuning functions are unchanged following apical dendrite ablation. **(a)** *Top:* 3D reconstruction of an example neuron before and after apical dendrite ablation. *Bottom:* Heat map depictions of the neuron’s single trial somatic calcium responses. Each column depicts responses to 12 directions (far left) before (D0) and 1 (D1), 3 (D3), and 5 days (D5) after ablation. Stimulus onset and offset are marked with broken yellow lines. Right insets: Zoomed-in view of the neuron’s responses to a rightward moving grating in 30 trials at D0 (blue dot) and D5 (red dot). Blue line: stimulus duration. **(b)** Peak-normalized orientation-tuning curves of the example neuron at D0 (black), D1 (red), D3 (purple), D5 (blue) and overlay of D0-5. **(c)** Polar plots depicting un-normalized orientation-tuning curves at each time point. **d-f**, scatter plots of **(d)** orientation preference, **(e)** orientation tuning width and **(f)** orientation selectivity index (OSI) from von-mises fitted tuning curves of each neuron before and 5 days after ablation. Blue, ablated neurons (n=11). Insets in d-f are bar plots of changes in (d) orientation preference (absolute-value change), (e) tuning width and (f) OSI in absolute values. Error bars are standard error of mean. Black, control neurons (n=12). P values are from T-test. Kruskal-Wallis Outputs for (d) orientation preference is p=0.23, (e) tuning width: p=0.64 and (f) OSI: p=0.42

We confirmed that targeted dendrites were successfully removed by generating a custom virus co-expressing GCaMP6s with activity-independent stable red fluorophore mRuby (Supp. Fig. 2). The elimination of the apical tuft following ablation was clearly identified with red fluorescent signal, and complete structural overlap was observed between GCaMP6 and mRuby. Ablation was further confirmed using post-hoc immunostaining with anti-GFP antibodies (Supp. Fig. 3). Post-hoc immunostaining of unlabeled axons and dendrites adjacent to the ablation site revealed no overt signs of damage (Supp. Fig. 3e, Supp. Movie 3) while adjacent GCaMP-labeled processes imaged *in vivo* near the ablation site were not noticeably affected (Supp. Fig. 1c), suggesting that ablation effects are spatially restricted within a radius of ~5 μm around the ablation point, in line with previous reports^14,15^.

### Loss of apical dendritic input does not alter orientation preference

L2/3 pyramidal neurons receive orientation tuned inputs scattered pseudo-randomly on their apical dendritic-trees^3,4,11,18^, but it is not clear whether these contribute to the neuron’s orientation preference. We tested the collective contribution of these inputs to orientation selectivity by ablating the apical tuft from L2/3 pyramidal neurons in fentanyl-dexmedetomidine sedated mice. The ablation of apical dendrite removes ~40% of the total excitatory input received by the neuron^19^. Figure 1c (top, Supp. Movie 2) shows the reconstructed morphology of one neuron before and after apical tuft ablation. Figure 2a illustrates single-trial dF/F responses of an example neuron to oriented gratings moving in 12 different directions, before as well as 1, 3 and 5 days after apical tuft ablation. Remarkably, removal of the entire apical tuft did not affect the neuron’s preferred orientation (Fig. 2b). The ablated neuron still responded maximally in response to its pre-ablation preferred orientation, and responses to other orientations were also very similar to pre-ablation (Fig. 2c).

Plotting the absolute-value change in orientation preference before and 5 days after ablation demonstrated that orientation preference did not change significantly following apical tuft ablation (Fig. 2d: blue circles corresponding to 11 ablated neurons all fall close to the diagonal as do black squares corresponding to 12 neighboring control neurons imaged together with ablated neurons). There was no significant difference in orientation preference change between the two groups (Fig. 2d, inset; p=0.13, T-test; p=0.23, χ^2^(1,21)=1.37, Kruskal-Wallis). Tuning-width (width at half maximum) and orientation selectivity index (OSI =[Response_pref_ -Response_null_]/[Response_pref_+Response_null_]) were also not significantly affected on average by apical tuft ablation (Fig. 2e, p=0.6; Fig. 2f, p=0.92; T-test). Pre-and post-ablation tuning curves for all recorded neurons are shown in Supp. Fig. 7. Bootstrap analysis to assess per-neuron shift in orientation preference and tuning width (see methods) also suggests that there is no consistent shift in orientation preference after removal of the apical dendrite beyond that observed in controls (Supp. Fig. 4–5, Supp. Table 1). Narrow distributions of orientation preference bootstrap estimates after ablation indicates that tuning of neurons remains robust even following dendrite ablation (Supp. Fig.4). No consistent changes in response amplitude or baseline firing rates were observed across neurons a day or more following ablation (Supp. Fig. 6, see figure legend for statistics). This demonstrates that, under the conditions tested, inputs to the basal dendrites are sufficient for determining the orientation preference and basic tuning function properties of V1 L2/3 pyramidal neurons^20^.

### Robustness of orientation-tuning to removal of multiple basal dendrites

We then applied our ablation strategy to the basal dendrites to explore how individual primary basal dendrites contribute to orientation selectivity (Fig. 3). Mouse V1 L2/3 pyramidal neurons typically have 5-8 (median 6) primary basal dendrites^19^. The point scan was targeted to a randomly-chosen primary basal dendrite, 5-10 μm away from the soma, before secondary branching. Neurons still present 24 hours post-basal dendrite ablation survived long-term and were studied. Overall, 36% of functionally studied neurons survived basal dendrite ablation (13 out of 33 neurons survived single basal dendrite ablation and 14 out of 41 neurons survived double basal dendrite ablation, from 30 mice). This was not significantly different from the survival rate after apical dendrite ablation (42%; p<0.01, p=0.51, Chi-square statistics = 3.29).

**Figure 3.**
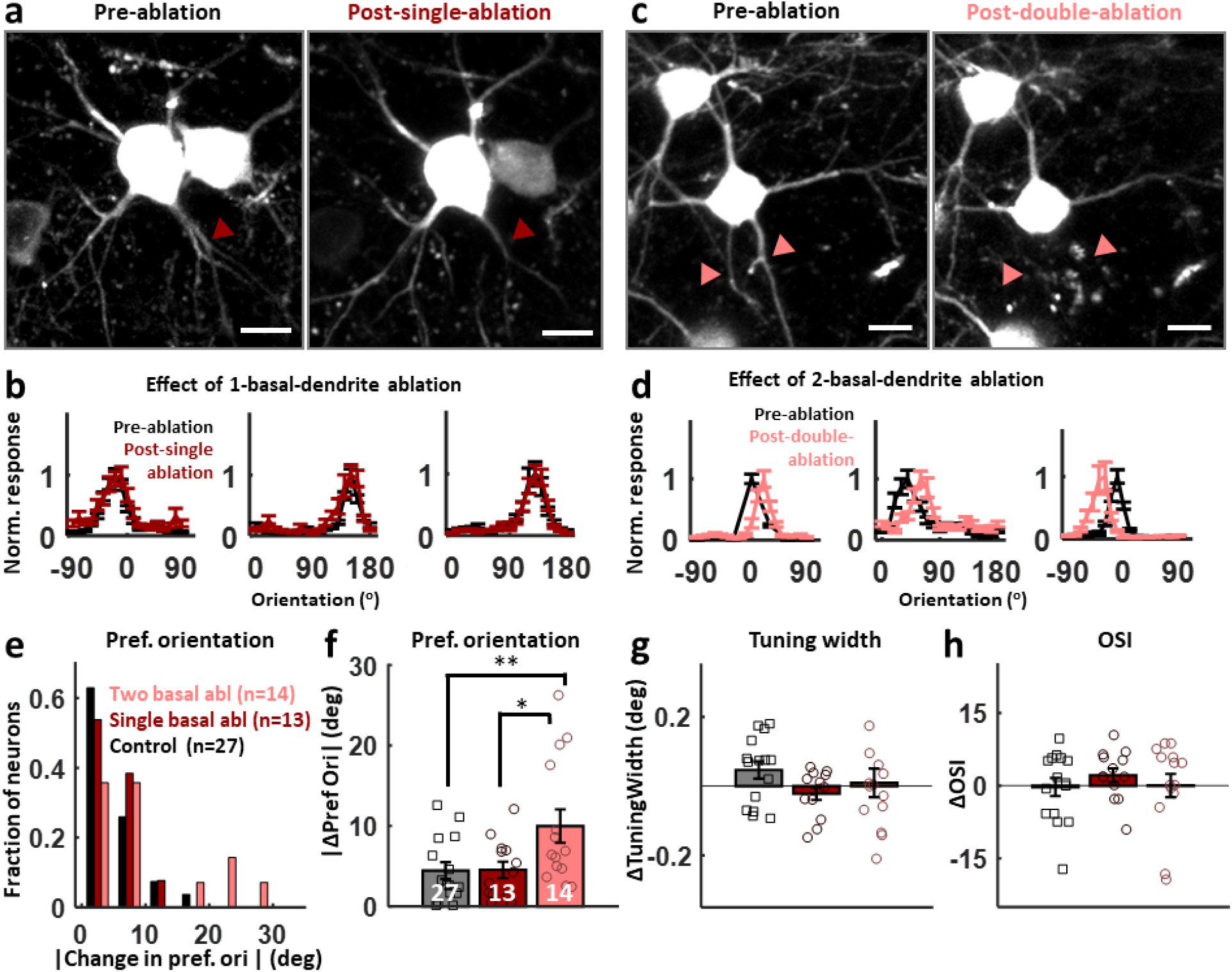
Removal of two, but not one, primary basal dendrites shifts orientation preference. **(a)** Z-stacked top view of an example ablated neuron and a neighboring control neuron before (left) and after (right) a single primary basal dendrite ablation. Ablated dendrite is marked with dark-red. **(b)** Three example peak-normalized orientation-tuning curves before (black) and 5 days after 1-basal-dendrite-ablation (dark-red). **c-d**, Figures in the same format with a-b for two primary basal dendrite ablation (marked with pink). **(e)** Histogram of the absolute-value change in preferred orientation (degrees) for **control (black, n=27), one (dark-red, n=13), and two (pink, n=14)** primary basal dendrites ablated neurons. **f-h**, Mean±SE of change in **(f)** preferred orientation (absolute-value changes), **(g)** tuning width (ANOVA output: F(2,51)=0.68, p=0.51, Kruskall-Wallis output: F(2,51)=0.53, p=0.77) and **(h)** orientation selectivity index (OSI, ANOVA output: F(2,51)=0.21 p=0.81, Kruskall-Wallis output: F(2,51)=0.33, p=0.84) from von-mises fitted tuning curves of each neuron 5 days after ablation. See Supp. Fig. 8 for histogram distribution of changes in tuning width and OSI. For (f), ** p=0.01, * p=0.03, ANOVA followed by Tukey test for multiple comparisons. ANOVA output: F(2,51)=5.1, p=0.0098. Kruskal-Wallis output: χ^2^(2,51)=6.48, p=0.039. For the control versus the 2-basal dendrite ablated groups p=0.045. Scale bars for (b) and (d) are 10μm.

Similar to what we observed with apical dendrite ablation (Fig. 2), cutting one basal dendrite (Fig. 3a, 8 to 12% of total dendritic input) did not alter the target neuron’s preferred orientation, nor its tuning-width or OSI (Fig. 3b, Supp. Fig. 7b). Neurons in which two basal dendrites were removed (Fig. 3 c, 16-24% of total input) also demonstrated remarkable tuning curve stability (Fig. 3d, Supp. Fig. 7b), but did show a small, yet significant, shift of ~12.5±2.7° on average (max: 30°) in orientation preference following ablation (Fig. 3e-f, p=0.0098, ANOVA with Tukey correction for multiple comparisons; Kruskal-Wallis output with Tukey correction for multiple comparisons: p=0.045 for the comparison control vs 2-basal; see figure legends for statistics). We again performed bootstrap analysis to estimate the reliability of orientation preference measurements neuron by neuron. The distribution of the bootstrap estimates of orientation preference before and after ablation was narrow, suggesting that the tuning properties of ablated neurons were measured reliably (Supp. Fig. 4–5, Supp. Table 1). Per-neuron bootstrap analysis also demonstrated that the number of two-basal dendrite ablated neurons that shifted their orientation preference post ablation was significantly different compared to controls (p<0.01; chi-square test), while this was not true for apical or single-basal dendrite ablations, which were not significantly different from controls. Tuning-width and OSI differences did not reach significance (Fig. 3g-h, p=0.51 (tuning-width), p=0.81 (OSI), ANOVA with Tukey multiple-comparisons test; See Supp. Figure 8 for histogram distribution of changes in tuning width and OSI). In one case, three (out of initially 7) primary basal dendrites were successfully ablated from one neuron, and even this cell maintained its orientation-tuning curve to a remarkable extent (just 10° shift following removal of 3 basal dendrites, Supp. Fig. 9a).

To test the possibility that the shift in orientation preference following 2-basal-dendrite ablation might be caused by a shift towards the tuning of the apical tree, we ablated the apical dendrite following 2-basal-dendrite ablation. This was successfully performed in two neurons, for which apical ablation did not further modify orientation preference (Supp. Fig. 9b,c and Supp. Movie 4). Thus, the orientation shift caused by 2-basal-dendrite cuts is unlikely to be due to a shift in the balance towards the apical tree’s orientation preference. This result further emphasizes the subordinate role that the apical dendrite plays in determining the orientation-tuning function of V1 L2/3 pyramidal neurons under the conditions of our experiment.

### Model-based exploration of tuning robustness following dendrite ablation

#### Apical dendrite ablation

Our finding that apical dendrite ablation has no effect on the orientation-tuning curve of L2/3 pyramidal neurons suggests a redundant contribution of the apical compared to the basal tree in orientation encoding. Otherwise, ablation should cause a change in neuronal orientation preference. To explore the possible input structures producing this result, we simulated a morphologically detailed L2/3 pyramidal neuron of mouse V1, with validated active and passive properties^6,21^ as well as literature-based synaptic density and single-synapse orientation-tuning properties (Fig. 4a-c and Supp. Fig. 10, see Supplementary Data for details of the model neuron)^4,19,22^. Three main parameters were varied: standard deviation (**σ_apical_**) of the distribution of single-synapse orientation preferences (pref_syn_) in the apical dendritic-tree, standard deviation (**σ_basal_**) of the distribution of pref_syn_ in the basal dendritic-tree, and the difference in the mean orientation preference of these two distributions (**Δ(μ_basal_, μ_apical_**)=|**μ_basal_-μ_apical_**|) (Fig. 4a, Supp. Fig. 10). We found that to generate tuning curves comparable to experiments (OSI≥0.2, tuning-width≤80°), (**σ_basal_, σ_apical_**) cannot be simultaneously larger than 45° (Supp. Fig. 11a-e). Interestingly, for all Δ(**μ_basal_, μ_apical_**) ranging from 0° to 90°, the model neuron’s orientation preference was consistently biased towards that of the basal tree (Fig. 4d), indicating the relative dominance of the basal dendrites.

**Figure 4.**
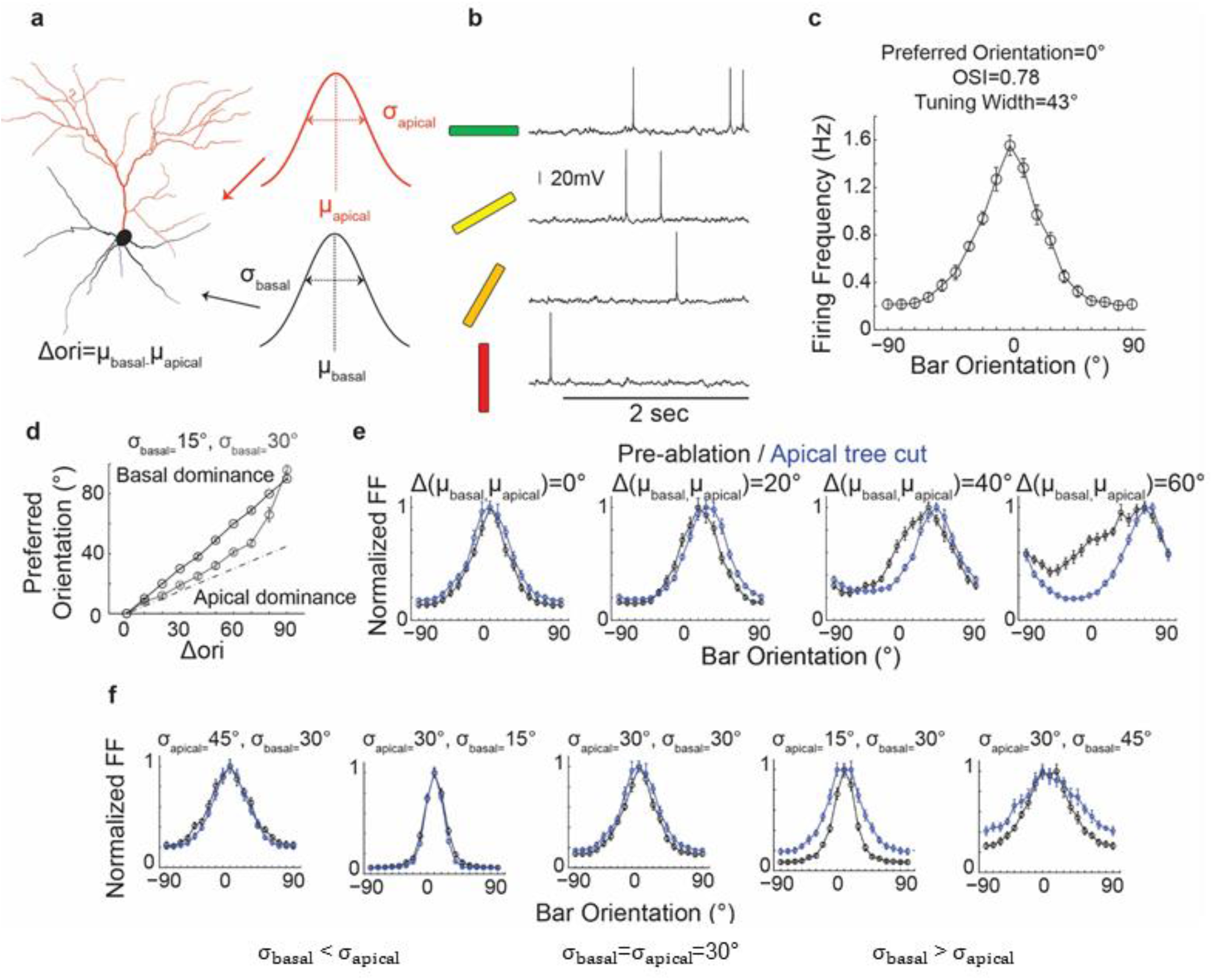
Simulations in biophysically-realistic model neurons suggest that apical and basal dendrites must have similar orientation preference to reproduce effects of ablation. **(a)** Reconstruction of the L2/3 pyramidal neuron used for simulation. Three main parameters, standard deviation (**σ**apical****) of the distribution of pref_syn_ in apical dendrites, standard deviation (**σ_basal_**) and mean (**μ_basal_**) of the distribution of pref_syn_ in basal dendrites, were varied. **μ_apical_** is arbitrarily fixed at zero. **(b)** The model neuron generates realistic subthreshold and action potential activity in response to 0°, 30°, 60° and 90° (left) orientations. Horizontal line represents stimulus duration (2s). (c) The model neuron displays realistic tuning curves (average tuning curve of 10 simulations when **σ_apical_= σ_basal_=30°**, **μ_apical_=μ_basal_ =0°**). **(d)** The model neuron’s preferred orientation is biased to the basal dendrites’ mean orientation preference for all Δ(μ_apical_,μ_basal_). Black line, **σ_basal_=15°**. Grey line, **σ_basal_=30°**. For both conditions, **σ_apical_=30°**. Dotted line indicates equal contribution of apical and basal trees in determining somatic orientation preference. (e) Left two panels: At low **Δ(μ_basal_, μ_apical_**) (<30°), ablation of the apical dendrite does not affect orientation tuning curve shape; Right two panels: At higher **Δ(μ_basal_, μ_apical_)** (>30°), apical dendrite ablation leads to a shift in orientation preference and narrowing of tuning width. **σ_apical_=σ_basal_**=30°. **(f)** Left two panels and the center: At **σ_apical_ > σ_basal_** or **σ_apical_=σ_basal_=30°**, ablation of the apical dendrite does not affect the tuning width; Right two panels: At **σ_apical_ < σ_basal_**, apical dendrite ablation broadens the tuning curve. **Δ(μ_basal_, μ_apical_)=0°**.

To further constrain the input parameter space, we modeled the effects of apical dendrite ablation on preferred orientation, OSI and tuning-width (Fig. 4e, Supp. Fig. 11f). We set **σ_apical_** at 30° as approximated from prior measurements of single-synapse orientation preference distribution on apical tuft dendrites (Fig. 4e, Supp. Fig. 11f)^11^. Figure 4e shows example tuning curves for **σ_apical_=σ_basal_=30°** with varying **Δ(μ_basal_, μ_apical_)**. For large **Δ(μbasal, μapical) (>40°**, Fig. 4e right, Supp. Fig. 11f upper half of heat-maps), the apical dendrite input functions as noise, broadening the orientation-tuning curve, and therefore its removal leads to shifts in orientation preference, narrower tuning-width and higher OSI, contrary to experiment. For intermediate Δ (20°-40°) and **σ_basal_** ≤ **σ_apical_** (30°), the basal dendrite dominates, and apical dendrite ablation has no effect (Supp. Fig. 11f), consistent with experiment. For **σ_basal_** > **σ_apical_** (30°), the apical dendrite input sharpens orientation-tuning, and therefore ablation of the apical dendrite leads to wider tuning width and lower OSI (Fig.5f right two panels, Supp. Fig. 11f). For small **Δ(μ_basal_, μ_apical_**) (0-10°, Fig. 4e left panel, Supp. Fig. 11f lower half of heat-maps), the model neuron’s preferred orientation does not change (mode change <10°) following ablation as long as **σ_basal_** is less than 60°.

**Figure 5.**
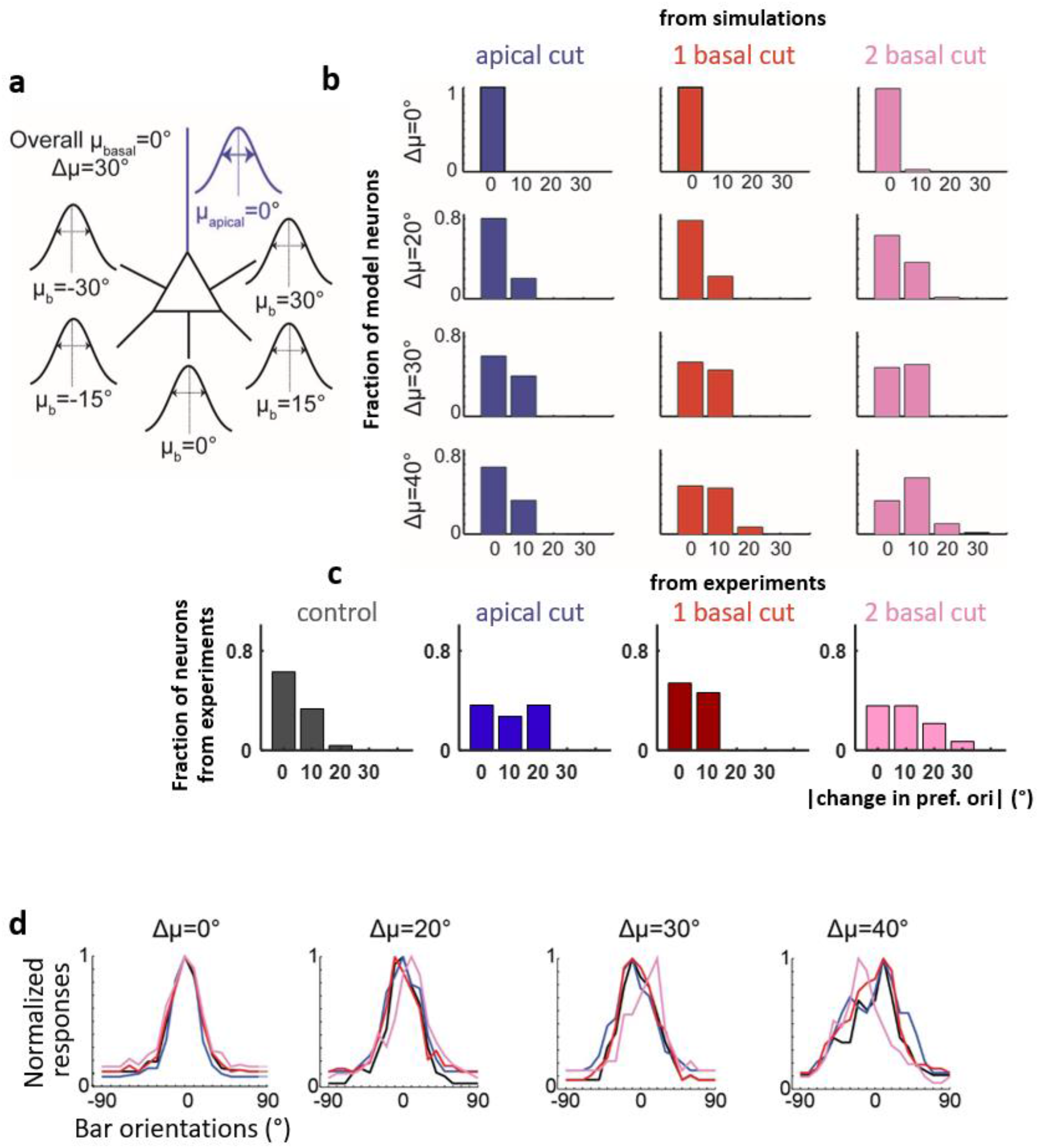
Simulated neurons with differentially tuned basal dendrites reproduce effects of ablation. **(a)** Example simulation schematic for disparity of orientation preference across basal dendrites, Δ_μ_=30°. μ_b_ for each basal dendrite was selected from a defined disparity μ_b_ ∈ {-**Δ_μ_ … μ_basal_ … Δ_μ_}**. Mean of pref_syn_ in the apical (**μ_apical_**) and basal (**μ_basal_**) were arbitrarily set to 0°. **σ_apical_=30°** and **σ_b_=σ_basal_=15°** for all basal dendrites. **(b)** Increasing disparity between basal dendrites (Δμ) leads to a mild shift in orientation preference with 2-basal ablation while minimally affecting tuning curves of apical or 1-basal ablation. Histogram of post-ablation absolute-value changes in orientation preference of model neurons with **Δμ=0°** (same as Figure 4), 20°, 30°, and 40° after apical (blue), single (red) and double (pink) basal dendrite ablation. For all, mean change in tuning-width is <10° and mean change in OSI is <0.2 (thresholds correspond to the mean+1std of the experimental data). **Δμ** of 50° or greater fail to generate tuning curves comparable to experiment (OSI≤0.2, tuning width≥80° for more than 30% of simulated neurons). **(c)** *In vivo* experimental data in the format of **(b)**, reproduced for comparison. **(d)** Example normalized tuning curves of model neurons with varying **Δμ** before (black) and after apical (blue), single (red) and double (pink) primary basal dendrite ablation.

#### Basal dendrite ablation: the shift vs. the drift hypothesis

There are at least two possible explanations for the basal dendrite ablation results in Figure 3. According to the **shift** hypothesis, the change in orientation preference following 2-basal-dendrite ablation is due to a shift toward apical tree inputs assuming **Δ(μ_basal_, μ_apical_)>0°**. Our exploratory observation that apical ablation following 2-basal-dendrite ablation does not shift the neuron’s orientation preference weighs against this hypothesis (Supp. Fig. 9b,c). Computational analysis, removing basal dendrites from model neurons with different **Δ** and **σ** (Supp. Fig. 11i,j), also failed to support the shift hypothesis. Although it was possible to generate model neurons with shifted orientation preference following 2-basal-dendrite ablation (i.e. for **Δ(μ_basal_, μ_apical_)**=30-50°, **σ_basal_**≤30°, **σ_apical_**=15°, mode change=10°, Supp. Fig. 11j), in these neurons apical ablation also led to a significant shift in orientation preference (Supp. Fig. 11h). There was no parameter set for which the 2-basal dendrite ablation, but not apical dendrite ablation, caused a shift in orientation preference.

The **drift** hypothesis entails that different basal dendrites are tuned to different preferences: cutting a subset of them will thus cause a drift due to loss of tuned basal inputs. To test this hypothesis, we generated model neurons with differentially tuned basal dendrites. The mean of the distribution of pref_syn_ in each primary basal dendrite (**μ_b_**) was set independently. **μ_b_** was selected from a range of **Δμ**, where **Δμ** represents the maximum deviation of one basal dendrite from the mean orientation preference of the soma (here arbitrarily set to 0°, Fig. 5a). **Δμ**=0° represents the null hypothesis, i.e. where all basal dendrites sample from the same salt-Q and-pepper arrangement of input orientation preferences centered at 0°. In this case, any postablation changes are due to sampling noise of the finite number of synapses and, as expected, basal dendrite ablation rarely causes changes in orientation preference (Fig. 5b, top row, Fig. 5d far left). The same was true for small disparity values (**Δμ**=20°, Fig. 5b second row, Fig. 5d). Intermediate disparity values (**Δμ**=30-40°) appear to be necessary to achieve a modest change (mode=10°, mean=8°, max=30° for Δ_μ_=40°) in orientation preference after 2-basal-dendrite cut, while maintaining stability of orientation preference following apical or 1-basal-dendrite cut, in agreement with experiment (Fig. 5b,c). Disparities ≥50° failed to generate tuning-curves compatible to the experimental data (OSI≤0.2 and/or tuning-width ≥80°). As expected, the neuron’s post-ablation changes in orientation preference strongly correlated with the magnitude of the change in mean orientation preference of the aggregates of the basal dendrites before and after two basal dendrite ablation (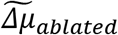, Supp. Fig. 12). The above (Fig. 5) were obtained using a narrow distribution of pref_syn_ in basal dendrites (**σ_b_= σ_basal_=15°)**. **σ_b_(=σ_basal_)** of 30° (Supp. Fig. 13) also induced a shift in orientation preference (max change=20°, mean=4.4°) but the fit to experimental data was worse than **σ_b_**=15°, indicating the need for sharply tuned input structures into basal dendrites.

In summary, these simulations suggest that to explain our experiment data, a) the difference in mean input tuning between apical and basal arbors, **Δ(μ_basal_, μ_apical_**), must be small, b) the distribution of input tuning to the basal dendrites should be similar to or narrower than the distribution of synaptic input to the apical tree (**σ_basal_ ≤ σ_apical_=30°**) and c) there needs to be a degree of heterogeneity of tuning (30°-40°) across basal dendrites.

## DISCUSSION

We showed that orientation selectivity in L2/3 pyramidal neurons, as measured by orientation preference, tuning-width and OSI, is robust to a complete loss of the apical tuft. This suggests that inputs to the basal dendritic domains are sufficient to determine the neuron’s orientation selectivity^20,23^. This is surprising given that apical dendrites receive orientation-tuned input ^11^, with at least four input sources accessible to apical dendrites that could potentially modulate orientation selectivity in V1: 1) feedback cortico-cortical projections from extrastriate cortex thought to refine orientation selectivity^9,10,24,25^, 2) direction-selective projections from the LGN^12^, 3) thalamo-cortical projections from lateral posterior thalamus^13^, and 4) dendrite-targeting inhibitory inputs^26,27^. Furthermore, several studies provided *in vivo* evidence of dendritic properties^28^ in V1 apical dendrites that have the capacity to contribute to the neuron’s orientation selectivity^6,29^, one arguing that apical tuft input functions to narrow the neuron’s tuning curve increasing selectivity^21^. However, these studies do not provide a causal connection between dendritic computations and the emergence of orientation selectivity at the soma. Our results suggest that with respect to orientation tuning apical dendritic tuft inputs are dominated by basal dendritic activity, at least under our experimental conditions. Moreover, apical dendrite ablation eliminates the contribution of regenerative potentials on distal dendrites^30^, which have been observed during complex behavioral task in vivo^31^ or L1 stimulation in vitro^32^. Our results suggest that neurons can still compute orientation tuning without those regenerative dendritic electrical events, favoring a feedforward model for orientation selectivity^33,34^. Computational modeling confirmed that basal dendritic input is dominant over a wide range of synaptic input parameters. It is interesting to explore in the future what behavioral conditions^35–37^ might change the relative contribution of apical dendritic inputs to orientation encoding in area V1.

Second, we showed that V1 neurons also remained robust to removal of 2-3 out of 5-8 primary basal dendrites. Specifically, only a small, though significant, change in orientation preference occurred in about half of the neurons that underwent two basal-dendrite ablation. The fact that we did not observe a marked loss of selectivity in any of the ablated neurons (Fig. 3) argues strongly against the possibility that there is a dominant, “master”, dendrite^17,38^, i.e. the situation in which input from a single dendrite dominates a neuron’s properties. Based on our results, we estimate the probability of a master dendrite dominated neuronal output to be p≈0.0003 for neurons with 6 basal dendrites (p≈(5/6)^13^x(4/6)^14^≈0.0003 corresponding to 13 neurons with single and 14 neurons with double basal dendrite ablation), or p≈0.0031 for neurons with 8 basal dendrites (p≈(7/8)^13^x(6/8)^14^≈0.0031).

Still, to explain the small but significant post-ablation change in orientation preference following 2-basal dendrite ablation, simulations suggest that orientation tuning is somewhat heterogeneous across basal dendrites, with basal dendrite orientation preference spanning a range of ~30°-40° (Fig. 5). Such heterogeneity may arise from dendrite-specific forms of plasticity potentially mediated by spatially-restricted biochemical signaling^39–41^ and/or dendritic spikes^6,28,29,42–44^.

These results give new insights into the complex structure-function relationship of the pyramidal neuron, the fundamental computational subunit of the neocortex. In particular, the remarkable robustness of orientation preference under dendritic micro-dissection hints at the extraordinary ability of sensory cortical neurons to maintain functional selectivity following input loss. Our approach emphasizes the importance of applying causal manipulations to study the contribution of dendritic arbors to sensory encoding. Dendritic microdissection is a powerful method for probing causal relations between dendritic structure/function and somatic properties that can be applied to several key questions in systems neuroscience research.

## Acknowledgments

Model neurons are available in ModelDB (accession number 231185) and its detailed description can be found in Supplementary Materials.

SS and JP were supported by NEI R01 grant EY-024019 and NINDS R21 grant NS088457. A.P. and P.P. were supported by the European Research Council Starting Grant dEMORY (GA 311435).

We thank Xiaolong Jiang for providing high quality Neurolucida reconstructions of L2/3 pyramidal neurons of mouse V1 visual cortex for the biophysical model study. We thank Matt Rasband for providing equipment and reagents for immunostaining experiments. We thank Sangkyun Lee for MATLAB expertise.

## Author contributions statements

J.P. and S.M.S. designed the study, A.P. and P.P. designed the modeling part of the study, J.P. and M.A.M. performed the experiments, J.P. analyzed the data, A.P. performed the simulations, J.P. and R.T.A. wrote the manuscript. J.P., R.T.A., A.P., P.P. and S.M.S. edited the manuscript together.

## Methods

All experimental protocols were approved by The Baylor College of Medicine (BCM) Institutional Review Board. Male and female wild-type (C57BL/6) mice were purchased from the institutional vivarium. Viral vectors were purchased from or edited/packaged by University of Pennsylvania Vector Core and BCM Vector Core.

### Sparse Labeling of L2/3 Pyramidal Neurons with GCamp6s, and Chronic Window Implantation

For viral injections and chronic window implantation, 6-10 week old wildtype C57BL/6 mice (both male and female) were anaesthetized with isoflurane (1-1.5%). Baytril (5mg/kg), Carprofen (5mg/kg) and Dexamethasone (1.5mg/kg) were administered subcutaneously. The depth of anesthesia was assessed via monitoring breathing rates. A headpost was implanted on the skull and a 3mm diameter craniotomy was made over the visual cortex of the left hemisphere. The craniotomy was centered 2.7mm lateral to the midline and 1.5mm posterior to the bregma. To sparsely label pyramidal neurons with GCaMP6s, a mix (~90 nl per site, up to 3 sites) of diluted CamKII-CRE (AAV5 or AAV1, diluted 40,000-120,000X) and flex-GCaMP6s (AAV5, diluted up to 2X) (U Penn Vector Core) or flex-mRuby-GCaMP6s (AAV8, BCM Vector Core) was injected slowly over 5 minutes per penetration using a Drummond Nanoject. Two to three penetrations ~0.5mm apart on average were performed per craniotomy. This approach enabled i) expression of GCamp6s at sufficient levels in each cell to image the dendritic tree, while ii) labeled neurons remained sparse. After the viral injection, a round coverslip was fitted to the craniotomy and sealed with vetbond and dental cement. Most chronic windows in our hands remain clear for 2-3 months. Visual stimulation and 2-photon imaging were performed on week 3-4 following viral injection, at which time GCamp6s expression was optimal.

### In vivo calcium imaging of sparsely labeled neurons

Three to four weeks after the injection, the GCaMP6s expressing mouse was sedated with Fentanyl (0.5 mg/kg) and Dexmedetomidine (0.5 mg/kg) (35) for imaging experiments. A stable level of anesthesia was confirmed by stable breathing rate and lack of movement. The right eye of the mouse was aligned to the center of the monitor. Customized light shielding was attached to the headpost to stimulate the mouse’s right eye effectively without producing light artifacts in the two-photon images. We imaged calcium activity of ablation candidate neurons and their neighbor control neurons in L2/3 of mouse primary visual cortex during visual presentation. Images were acquired at ~9 frames/sec using an Ultima IV microscope in spiral scanning mode with a 20X, 0.95 NA, Olympus objective or 25X 1.0 NA Nikon objective (5-30mW laser power, 900 nm). GCamp6s gives excellent signal to noise ratio, corresponds well to the underlying firing rate, and is well suited for measuring tuning functions. Orientation-tuning measurements and structure imaging were repeated before (day 0) and 1, 3, 5 days after ablation.

### In vivo dendrite ablation

Under two-photon scanning, fluorescent dendrites were clear and visible at low laser power (<20 mW, 910nm). Dendritic arbors of several L2/3 neurons were imaged using a custom Ultima IV 2-photon microscope. First, we screened for clearly orientation-tuned neurons with a clearly defined primary apical dendrite (Primary apical bifurcation >20 μm away from the soma, soma depth between 170 and 250 μm, average pre-ablation orientation selectivity index for all ablated neurons was 0.79±0.18, max=0.99, min=0.38). Target apical dendrites were severed at least 15 μm away from the soma (Fig. 1c and Supp. Movie 1, 2). Basal dendrites were severed at least 10 μm away from the soma (Fig. 3a,c and Supp. Movie 4).

The ablation point was magnified 13-15 times with 1024 resolution at 910 nm, then ablated via repeated 200-400ms point scans at 150-200mW power, 800 nm wavelength. Approximate area impacted by single point scan is an ellipsoid of 0.4 μm (x,y) and 1.2 μm (z) diameter according to our point spread function (Supp. Fig. 1a). Differences in ablation parameters depended on depth of ablation plane, overlying shadow casting vessels, and window clarity. As reported in previous studies (11, 12, 36), when a single point on a fluorescing dendrite received several focused laser pulses, the targeted dendrite formed a beads-on-a-string morphology immediately after the ablation and then degraded within a day (Supp. Fig. 1). Successfully ablated dendritic segments become transiently brightly fluorescent as calcium enters the membrane, then recover its original brightness. Distal to the ablation point, dendritic segments displayed marked beading within ~1 hr and degraded completely within a day. Dendritic segments proximal to the ablation point by about 5-10 μm survive indefinitely. About 70% of ablated neurons recovered to baseline fluorescence levels within 2-3 hours post-ablation. The remaining 30% degenerated and disappeared within 24 hours. For functionally mapped neurons, we found the similar chance of survival rate both from apical (11 out of 26, from 19 mice) and basal dendrite ablation (13 out of 33 for one basal dendrite ablation and 14 out of 41 two basal dendrite ablation, from 30 mice). *χ*^2^ statistic for the survival rate of apical and basal cut experiments is 3.29. The p-value is 0.51. The result is not significant at p < 0.01. Surviving neurons did not demonstrate significant regrowth or degradation at the ablation point (at most 3-5 μm regression toward the soma). The remaining dendritic arbor, as well as neighboring neuronal structures, maintained their original structure for at least 14 days following ablation (Fig. 1b, Supp. Fig. 1b).

We assessed the possibility of injury to the neuron and/or neuropil in four ways. Evidence for minimal post-ablation damage includes: 1) ablated neurons survived indefinitely after being stressed for a while (as documented by the calcium influx and increase in fluorescence, Supp. Fig. 1d), 2) no visible change in spontaneous or visual-evoked calcium activity were observed in the days following ablation of either apical or basal dendrites (Fig. 1–3, Supp. Fig. 4,5), 3) no morphological changes were observed in non-ablated structures from the same neuron or processes of nearby non-ablated neurons, even 5 microns from the ablation site (Supp. Fig. 1), and 4) immuno-labeled neuronal processes (anti-Tuj1) were minimally affected at the ablation site (Supp. Fig. 3 and Supp. Movie 3). Other labs using the same method have shown no effect of this ablation on spine morphology or turnover in remnant proximal segments of ablated dendrites (36), and FIB-SEM reconstructions of ablation sites found a ~5 micron lesion with no prominent glial scar 5 days post-ablation (12). Taking all of the data together leads us to conclude, as others have (11, 12, 36), that the off-target damage mediated by 2-photon microdissection is minimal.

### Visual stimulation

Visual stimuli were generated with MATLAB (Mathworks Inc.) PsychToolBox and presented on a Dell monitor (77°x55° of visual angle), at a fixed mean luminance (80 candela/m2), positioned 32 cm in front of the animal. Prior to orientation mapping the animal was adapted for at least 15 minutes to the mean luminance level. To measure orientation selectivity, grayscale square-wave gratings (0.04 cycles/degree, 2 cycles/second) moving in one of 12 (30° steps) or 36 (10° steps) directions were presented in pseudorandom order over the full stimulation field. Stimulus presentation lasted 2 sec and the inter-stimulus interval (uniform illumination set at the mean intensity) lasted 3 seconds. Twenty to thirty repetitions per stimulus direction of motion were collected and analyzed.

### Data Analysis

Calcium trace from each soma was selected and separated using custom MATLAB functions, including the ROI selecting function ‘roigui’ from T.W. Chen at Janelia Institute. Baseline fluorescence (F0) for calculating (F-F0)/F0 was the average of the 20% lowest values in a 20 sec (10 sec pre, 10 sec post) window around each frame. Visual-evoked calcium responses arose ~200ms after the onset of the stimulus and peaked around the offset of the stimulus. The per-trial response was calculated as the average of a 1.5 second window centered at the peak of the mean response across all orientations. Tuning curves were calculated by averaging responses across repetitions of each stimulus orientation. From this tuning curve, orientation selectivity index was calculated with (R_pref_-R_ortho_)/(R_pref_+R_ortho_). All orientation-tuning curves were fitted with von Mises function using Circstat tool box^45^ after baseline subtraction. Preferred orientation and tuning width was calculated from the fitted tuning curve. Calculating preferred orientation and tuning width from the raw mean tuning curve generated similar results. To assess the effect of dendrite ablation on tuning curves, preferred orientation, tuning width and OSI of before ablation values were subtracted from 5 days after ablation value of the corresponding neurons. We used both ANOVA and Kruskall-Wallis test to assess the significance between shift in ablated neurons and control neurons. Tukey test for multiple comparisons was used for basal dendrite ablation results. F statistics (for ANOVA) and Chi-square statistics (for Kruskall-Wallis) were provided together with p-values for comparing control, 1-basal and 2-basal dendrite ablation neurons. P-values between specific groups (e.g. control vs 2-basal dendrite ablation) are from the statistics with multiple comparisons. Absolute-value was used to assess the shift in preferred orientation.

For bootstrapping of Supp. Fig 4–5, five single-trial responses were randomly subselected from the 20-50 trials acquired per orientation to generate simulated tuning curves. The preferred orientation was calculated from the von Mises-fit orientation tuning curve based on the sub-sampled data. This was repeated 1000 times to generate confidence intervals for the estimate of the preferred orientation pre- and post-ablation per neuron. Distributions of the orientation estimates were shifted to have the mean of the preferred orientation estimates before ablation at 0°. We defined a neuron with significant difference in preferred orientation (or in tuning width) if 95% of confidence intervals of pre- and post-ablation do not overlap to each other. The frequency of detecting neurons with significant difference in preferred orientation pre- and post-ablation was assessed with chi-square test.

### Immunofluorescence labeling

Mice were killed by isoflurane overdose followed by cervical dislocation and decapitation. Following decapitation, brains were quickly removed and fixed in ice cold 4% PFA (pH 7.2, 1 hour) followed by 20% sucrose immersion for 24 hours and then 30% sucrose immersion for 24 hours. Fixed brains were frozen in OCT medium and sectioned into 50μm coronal sections. For immunostaining, free floating sections were washed in 0.1 M PB and blocked in PBTGS (0.1 M PB with 0.3% Triton X-100 detergent and 5% goat serum) for 1 hour. Sections were incubated in primary antibody over night at room temperature. The following antibodies were used: chicken anti-GFP (Abcam) and mouse anti-beta III Tubulin (Santa Cruz). Primary antibodies were then labeled using AlexaFluor-conjugated secondary antibodies (Invitrogen). Sections were visualized with a Zeiss Imager, Z1 fluorescent microscope with an Apotome attachment.

### Biophysical Model

The morphologically detailed L2/3 V1 pyramidal neuron model (Figure 4a) was implemented in the NEURON simulation environment ^46^. Model neurons (available in ModelDB, private model until publication, https://senselab.med.yale.edu/modeldb/enterCode.cshtml?model=231185, password: nassi123) were constrained against experimental electrophysiological and anatomical data^6,47–52^ (see Supplementary information for detailed description). Synapses impinging in the apical and basal dendrites where either spontaneously activated or stimulus driven. Stimulus driven synapses were assigned with preferred orientations and randomly distributed along the dendrites ^4^ When assigning orientation preference to synapses, we varied the standard deviation of the distributions (σ_basal_, σ_apical_), the difference (Δ) of μ_apical_ and μ_basal_, as well as the μ_b_ of each individual basal dendrite (Δμ). Control condition was the one corresponding to σ_apical_=σ_basal_=30°, μ_apical_=μ_basal_=0°, Δμ=0°, under which the OSI was 0.78 and the tuning width 43° (Fig. 4b,c). Ablation was simulated by removing the respective dendritic compartments and adjusting excitatory synaptic weights for the change in neuronal excitability. For each condition, we calculated the preferred orientation, tuning width and OSI of the resulting somatic tuning curves.

## Supplementary Information

**Supplementary Figure 1.**
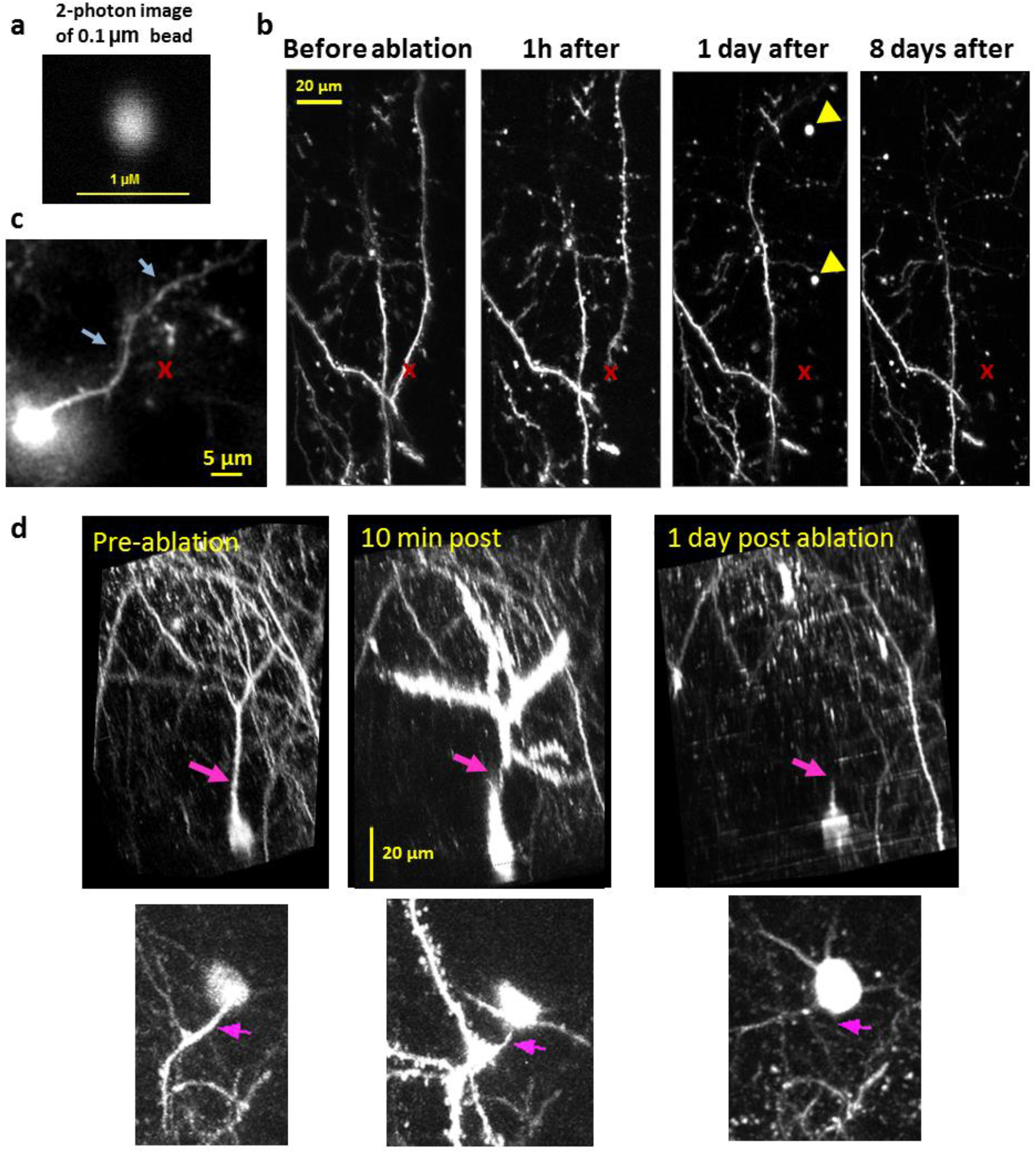
Visual confirmation of apical dendrite ablation. **(a)** 2-photon image of a 0.1 micron fluorescent bead, revealing the resolving power (and point scan size) of the microscope to be about 0.4 μm in the X-Y plane (~1.2 μm in Z). **(b)** 5-μm-thick maximum Z-projection of apical dendrites of a GFP expressing L2 pyramidal neuron imaged in vivo from a thy1-GFP mouse before and 1 hour, 1 day and 8 days after ablation. A point scan with 800nm wavelength, 200ms duration, and ~150 mW power was applied to the point marked with the red ‘x’. Dendritic segments distal to ablation point developed a beads-on-a-string appearance within 1 hour postablation, and disappeared by 24 hours. GFP-filled remnants of the ablated dendrite were visible 24 hours after ablation (yellow arrows), but disappeared by 8 days. Dendrites near the ablation point, even adjacent branches from the same neuron, showed no change in morphology post-ablation. **(c)** Single slice through an example apical ablation site (‘red x’), acquired 5-days post-ablation (target neuron tuning curve shown in Figure 2a-c). GCaMP-labeled dendrites from nearby control neurons did not show any fine-scale changes following ablation, even ~5 μm away from the ablation site (pale blue arrows). (d) Side-projection (top) and top-projection (bottom) of an example neuron before (left), 10 minutes after (middle) and 1 day after 2-photon ablation of the apical dendrite. Pink arrow indicates ablation point. Neuron increased its fluorescence levels immediately after ablation likely due to electrolyte influx through the instantaneous injury caused by ablation. By 1 day following ablation target neurons’ fluorescence has returned to baseline, and only a short remnant of the apical dendrite remains.

**Supplementary Figure 2.**
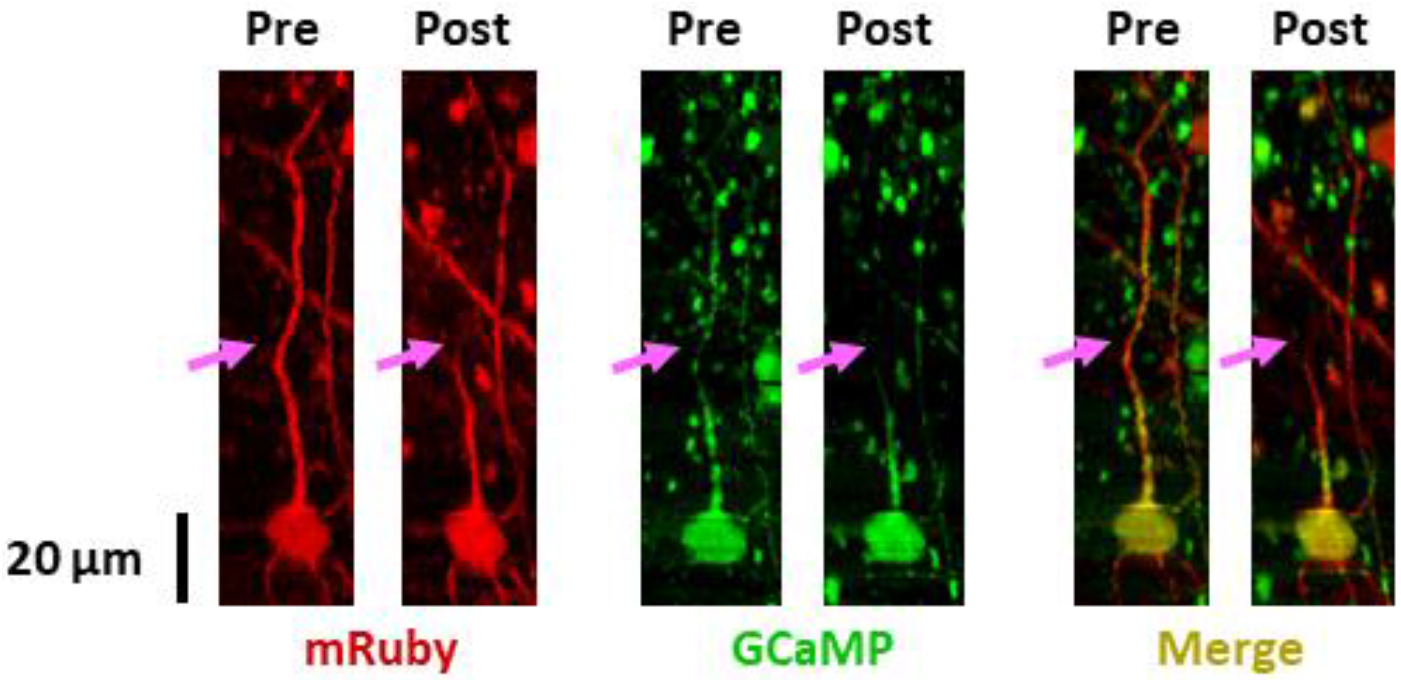
Confirmation of ablation by co-expression of red fluorophore mRuby with GCaMP. Custom engineered virus (AAV8-flex-mRuby-2A-GCaMP6s) was coinjected with low titer (1:80,000) AAV1-CaMK2-Cre to enable co-expression of an activity-independent red fluorophore with GCaMP6. Side-projection of target ablated neuron in the red channel (left), green channel (middle) and merged channels (right), showing overlap of GCaMP6 and mRuby expression in the neuron before and 5 days after ablation of the apical dendritic tuft (pink arrow depicts ablation point). Vertical variation in the green signal (middle panels) is due to fluctuations in GCaMP fluorescence which reflect activity of the cell during z stack acquisition.

**Supplementary Figure 3.**
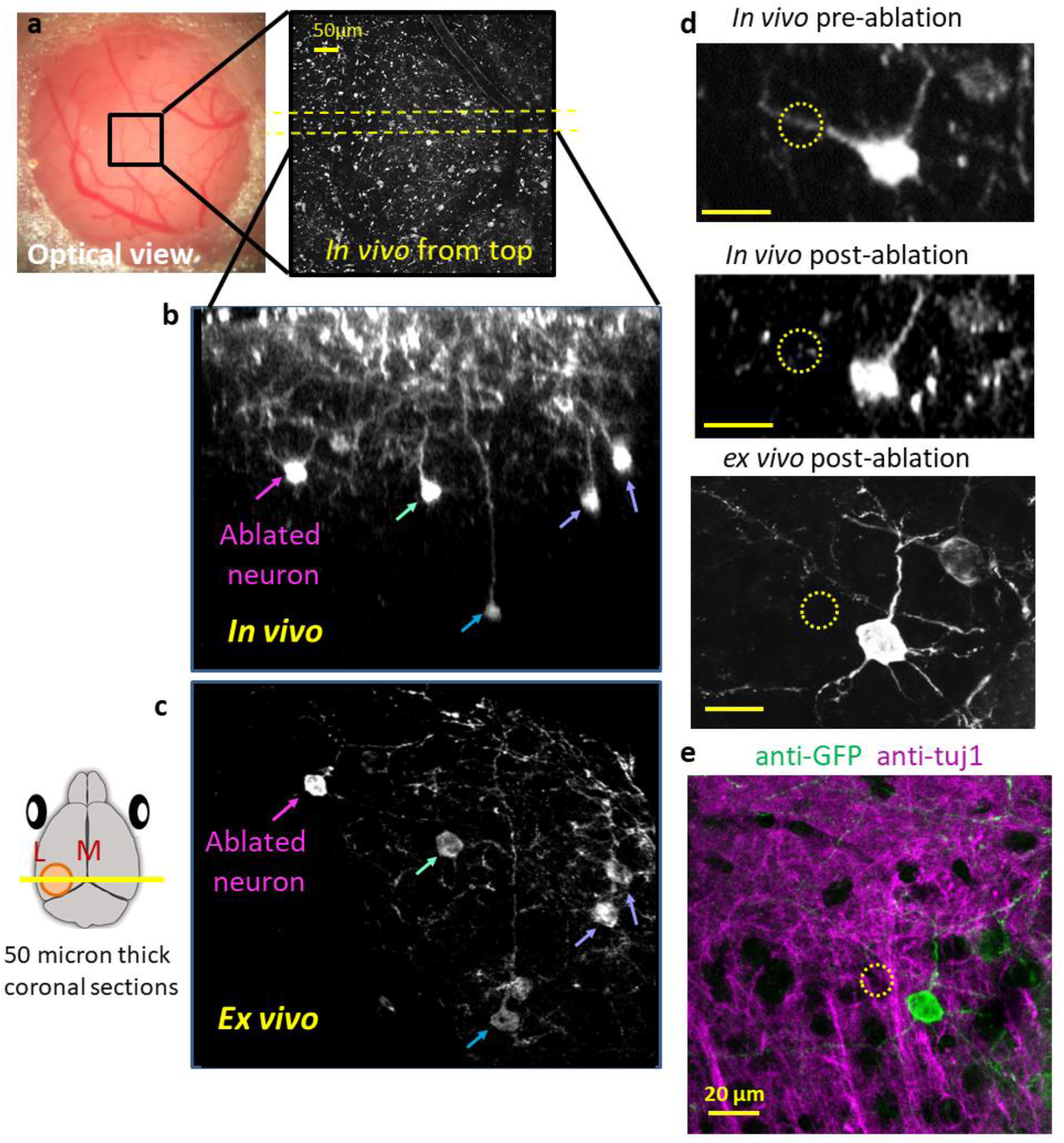
Post-hoc immunohistochemical confirmation of ablation and assessment of damage at ablation site. Dendrites of five neurons in this brain were successfully ablated. Cells were chosen at the edge of the ~400-micron-diameter sphere of GCaMP expression to ease locating them post-hoc. One day after ablation, the brain was fixed in paraformaldehyde, and 50-micron-thick coronal sections were made through the visual cortex, tracking each slice from posterior to anterior. Slices were immunostained with anti-GFP and anti-Tuj 1 antibodies to visualize GCaMP-labeled and unlabeled neuronal processes, respectively. **(a)** *<u>Left:</u>* Optical image of craniotomy and imaging window. *Right:* two-photon z projection of the boxed region from left panel. Note the same surface blood vessels are visible in each image (arrows identify corresponding blood vessels). **(b)** *<u>In vivo</u>* coronal side view of an ablated neuron (pink arrow) and identified neighboring labeled neurons (other arrows). **(c)** *Ex vivo* image of same neurons visualized in vivo in panel b. **(d)** Zoomed-in views of the ablated neuron before ablation in vivo (top) after ablation in vivo (middle) and ex vivo (bottom). Note the similar orientations of the remaining dendrites emanating from the soma, as well as the location of a neighboring neuron. Yellow circle centered at ablation site. Diameter of the yellow circle is 12 micron in (d) and (e). All scale bars in (d) and (e) are 20 microns. Images acquired in vivo (top two panels) were 3D rendered and rotated to optimally display the ablated dendrite. Apparent differences in structure sizes are due to differences in viewing angle. **(e)** Immunohistochemical assessment of damage at ablation site. Green shows anti-GFP signal (amplifying GCaMP), magenta shows anti-Tuj1 signal (antibody specific to axonal and dendritic microtubules). No clear sign of disturbed processes were observed around the ablation site as confirmed previously with EM (See Fig. 2 of ^12^). Note that the black regions that appear to be within the dotted circle (ablation target) do not actually represent lesions but unstained regions corresponding to cell bodies appreciated better on a different plane. These are more clearly represented in animated Z-stacks, Supplementary Movie 3.

**Supplementary Figure 4.**
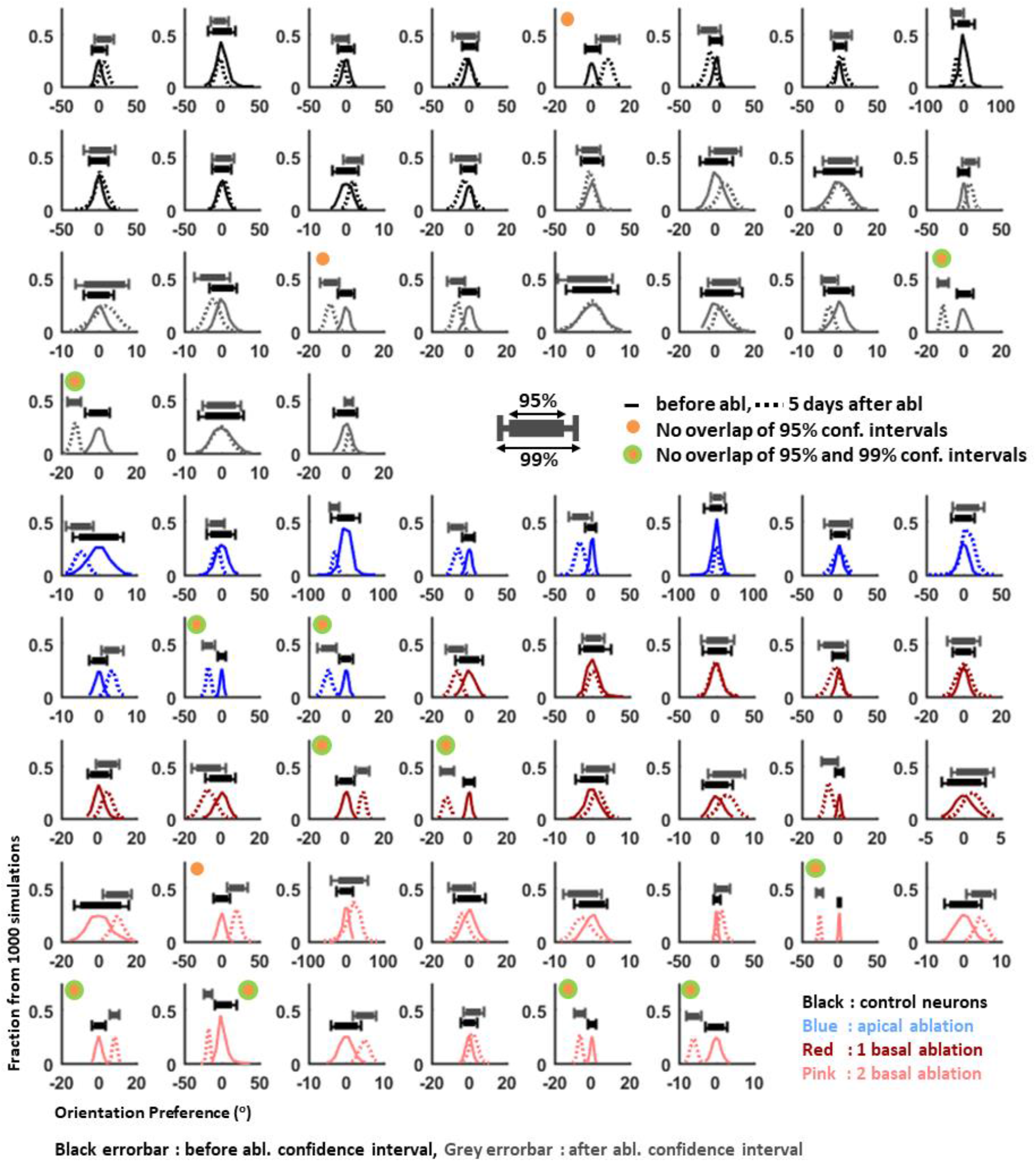
Distribution of bootstrap estimates of neurons’ orientation preferences before and after ablation. Histogram of the distribution of bootstrapped estimates of the orientation preferences before (solid line) and after ablation (dotted line) for neurons with apical dendrite (blue, n=11), 1 basal dendrite (dark red, n=13), 2 basal dendrite (pink, n=14) ablation and control neurons (black, n=27) from 1000 simulations. Five single-trial responses were randomly sub-selected from the 20-50 trials acquired per orientation to generate simulated tuning curves. The preferred orientation was calculated from the von Mises-fit orientation tuning curve based on the subsampled data. This was repeated 1000 times to generate confidence intervals for the estimate of the preferred orientation pre and post ablation per neuron. Note that all neurons have a narrow distribution for the orientation preference even after ablation, which indicates highly robust tuning even after dendrite ablation. Distributions of the orientation estimates were shifted to have the mean of the preferred orientation estimates before ablation at 0°. Errorbars above the histograms indicate the 95% (filled box) and 99% (tick to tick) confidence intervals before (black) and after (grey) ablation. A yellow dot indicates that there is no overlap between the 95% confidence intervals of the pre-and post-ablation orientation preference estimates. Green ring indicates that there is no overlap between 99% confidence intervals of the pre- and post-ablation orientation preference estimates. 6 out of 14 two-basal-dendrite-ablated neurons showed significant separation of the distribution of orientation preference estimates before and after ablation (yellow dot) while 2 out of 13 one-basal-dendrite-ablated, 2 out of 11 apical-dendrite-ablated, and 4 out of 27 control neurons did. The frequency of detecting neurons with significant difference in preferred orientation pre- and post-ablation was significantly different than controls as assessed by the χ^2^ statistic. Specifically, for 2 basal vs control is χ^2^=3.9 (p=0.04), while p was not significant for either 1-basal (p=0.96) or apical dendrite ablation (p=0.79) versus controls.

**Supplementary Table 1.** Chi-square test comparing control and ablated neurons for finding neurons with significant shift in preferred orientation.

| Chi-square test comparing the chance of having neurons with significant shift in orientation preference | Apical vs Control<br>p-value ($\chi^2$) | 1Basal vs Control<br>p-value ($\chi^2$) | 2Basal vs Control<br>p-value ($\chi^2$) |
| --- | --- | --- | --- |
| (95% confidence interval) | 0.79 (0.07) | 0.96 (0.002) | **0.04 (3.9)** |
| (99% confidence interval) | 0.33 (0.96) | 0.43 (0.62) | **0.02 (5.2)** |

**Supplementary Figure 5.**
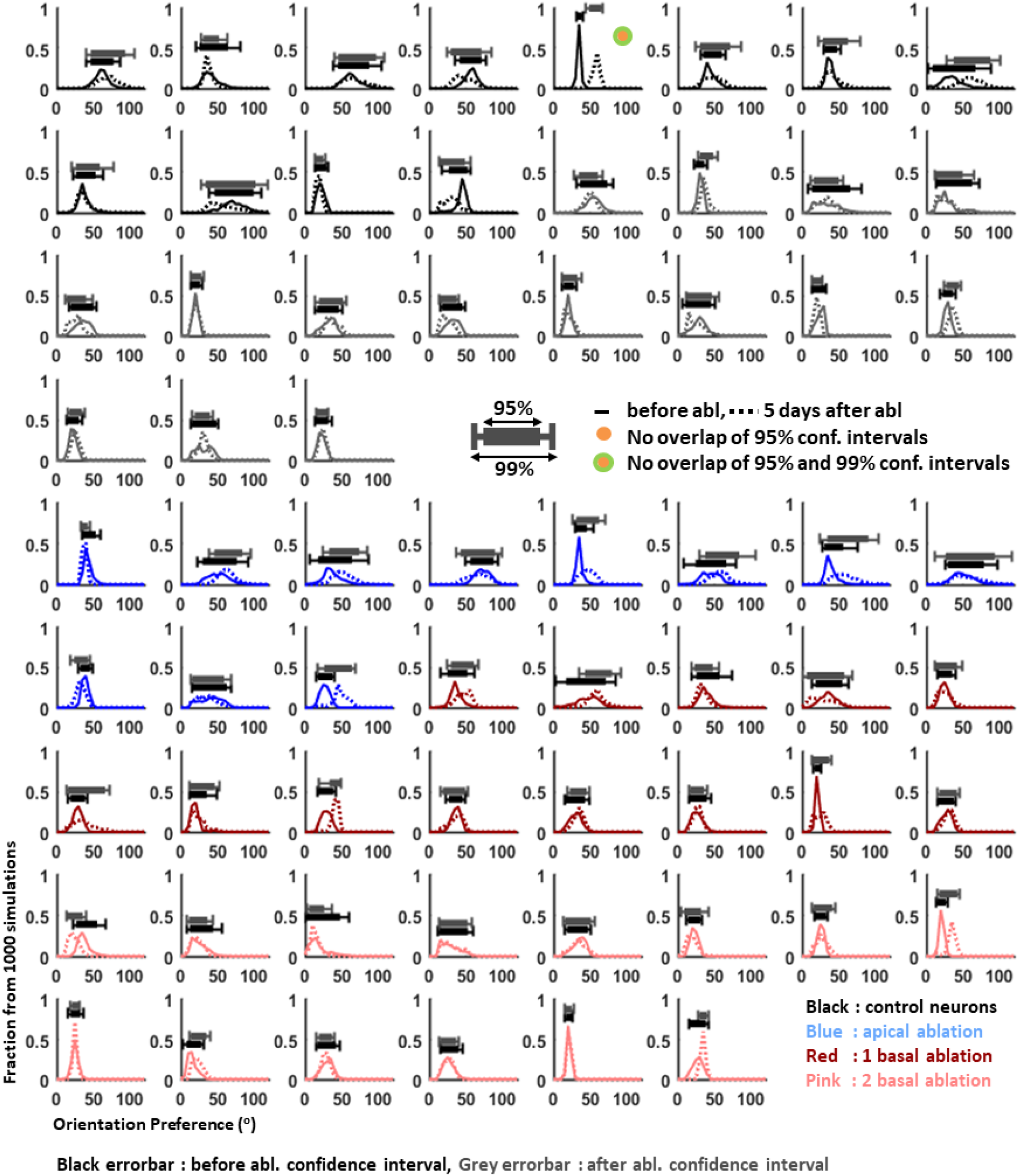
Distribution of bootstrap estimates of neurons’ tuning width before and after ablation. Histogram of the distribution of bootstrapped estimates of tuning width before (solid line) and after ablation (dotted line) for neurons with apical dendrite (blue, n=11), 1 basal dendrite (dark red, n=13), 2 basal dendrite (pink, n=14) ablation and control neurons (black, n=27) from 1000 simulations. Five single-trial responses were randomly subselected from the 20-50 trials per orientation to generate simulated tuning curves. The tuning width was calculated from mean orientation tuning curve based on the sub-sampled data. This was repeated 1000 times to generate bootstrap confidence intervals for the estimate of the tuning width pre- and post-ablation per neuron. Errorbars above the histograms indicate the 95% (filled box) and 99% (tick to tick) confidence intervals before (black) and after (grey) ablation. A yellow dot indicates that there is no overlap between the 95% confidence intervals of the pre-and post-ablation tuning width estimates. Green ring indicates that there is no overlap between 99% confidence intervals of the pre- and post-ablation tuning width estimates. Only 1 out of 54 neurons (in the control condition) showed a significant shift in tuning width by this measure.

**Supplementary Figure 6.**
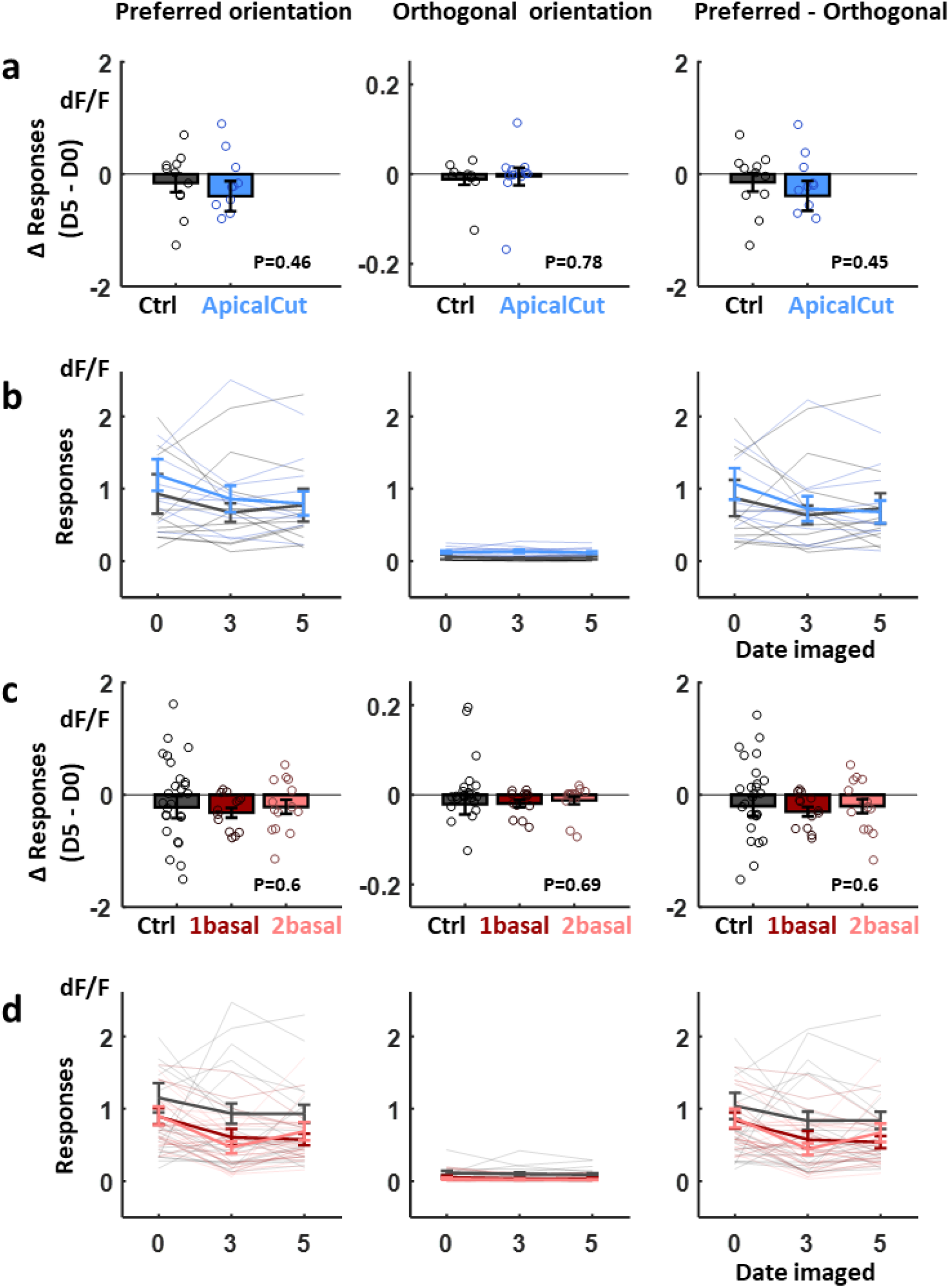
Response amplitude across days in ablated neurons are comparable to those of control neurons. **a**, mean±SEM of changes in responses to preferred orientation (left), orthogonal orientation (center) and gain [(response to preferred orientation) – (response to orthogonal orientation), right] before and 5 days after apical dendrite ablation. P-values are from Mann-Whitney U test. **b**, individual lines represent mean response of a single neuron from 20-30 trials in response to its preferred (left), orthogonal orientation (center), or the differences between preferred and orthogonal orientations (right). Day 0 corresponds to pre-ablation. Thick lines are averaged value across mean responses of individual neurons. Error bar is SEM across neurons. (a)-(b), blue: apical dendrite ablation, black/grey: control neurons imaged together with ablated neurons. **c**, mean±SEM of changes in responses to preferred orientation (left), orthogonal orientation (center) and gain [(response to preferred orientation) – (response to orthogonal orientation), right] before and 5 days after 1 basal and 2 basal dendrite ablation. P-values are from Kruskall-Wallis with Tukey test for multiple comparison. Outputs are χ^2^(2,51)=1, p=0.6 (preferred orientation, left), χ^2^(2,51)=0.7, p=0.69 (orthogonal orientation, center) and χ^2^(2,51)=1.02, p=0.6 (pref-ortho). **d**, figures for basal dendrite ablation in the same format with (b). (c)-(d), dark red: 1 basal dendrite ablation, pink: 2 basal dendrite ablation, black/grey: control neurons.

**Supplementary Figure 7.**
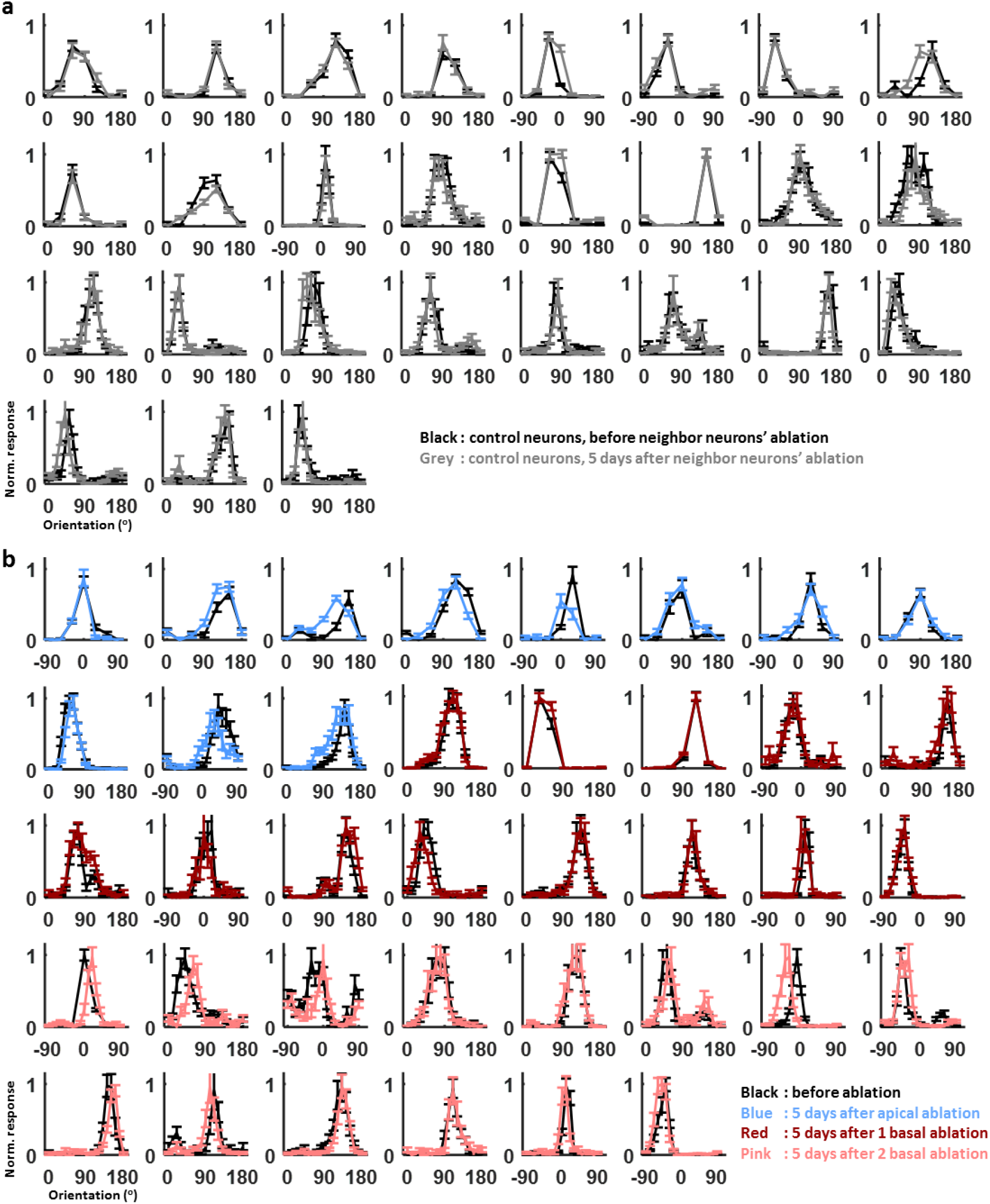
Peak normalized orientation-tuning curves of control and ablated neurons. All tuning curves in (a) and (b) were baseline-subtracted and peak-normalized for illustration purposes only. **(a)** Tuning curves of control neurons before (black) and 5 days after (grey) neighbor neurons’ ablation. **(b)** Tuning curves of ablated neurons. Pre-ablation tuning curves are in black. 5 days post-ablation tuning curves are in blue (apical dendrite ablation), red (single primary basal dendrite ablation) or pink (double primary basal dendrite ablation). Although there are occasional L2/3 pyramidal neurons that show small shifts in orientation preference following apical dendrite ablation (2/11), control neurons (4/27) or single basal dendrite ablation (2/13), only neurons with 2 basal dendrite ablations show significant orientation preference shift on average (10/14, ~12.5° on average, see Fig. 3f). Furthermore there was no significant change in tuning width or orientation selectivity index on average across each condition (Fig. 3g,h), although 4/11 neurons after apical dendrite ablation did show slightly widened tuning functions.

**Supplementary Figure 8.**
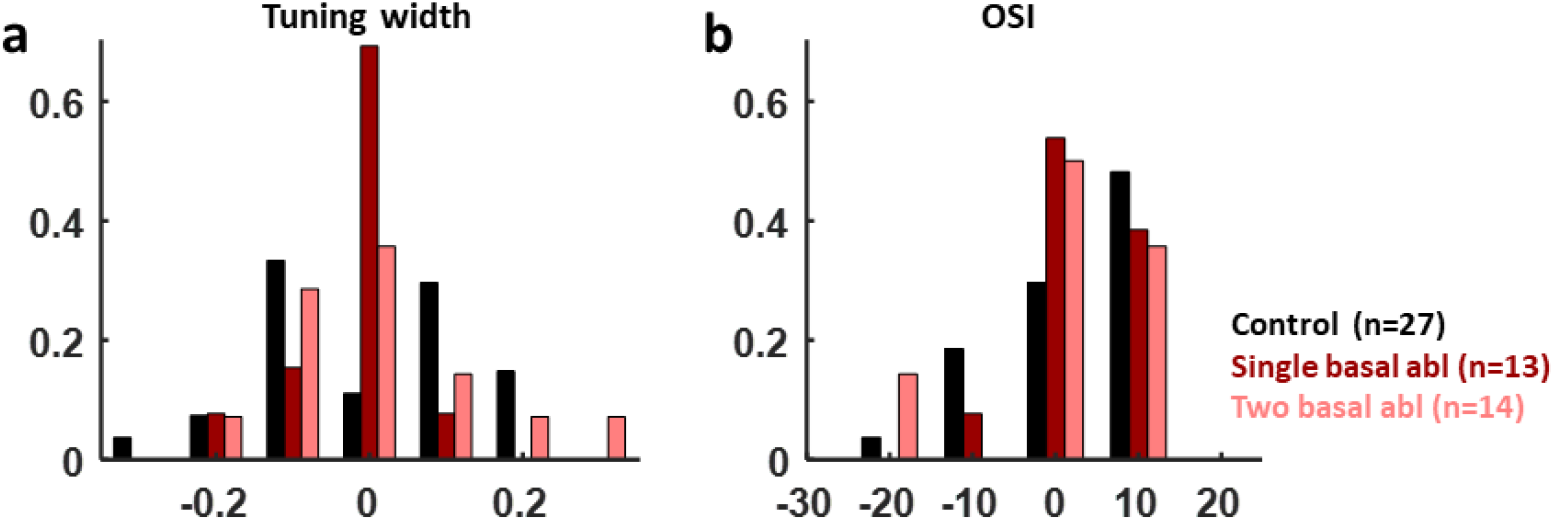
**Histogram of the change in tuning width** (a) and orientation selectivity index (b) for control (black, n=27), one (dark red, n=13), and two (pink, n=14) primary basal dendrites ablated neurons. Kruskall-Wallis output for tuning width (a): F(2,51)=1.29, p=0.28 and for OSI (b): F(2,51)=0.33, p=0.8).

**Supplementary Figure 9.**
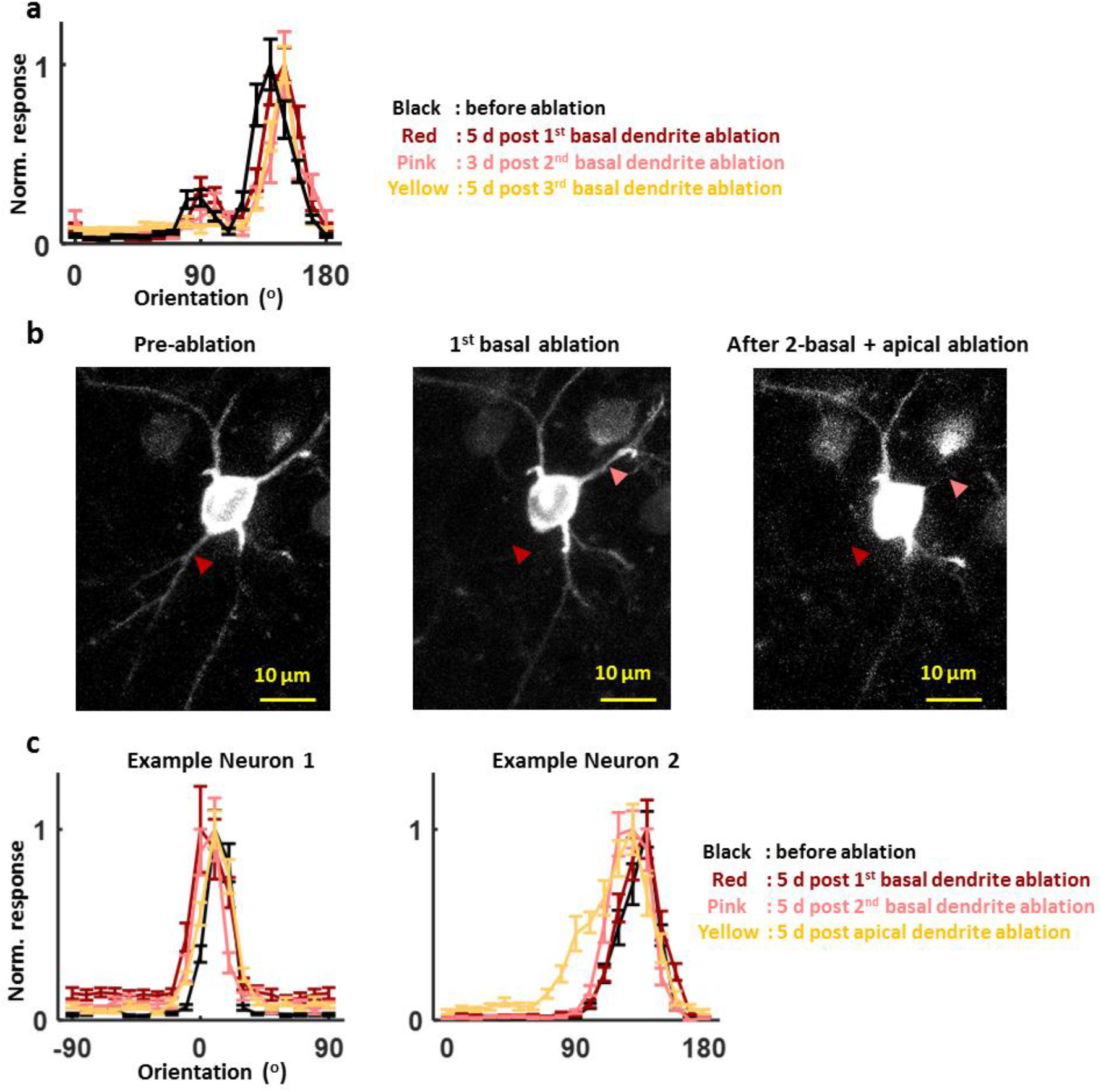
Stable orientation preference following multiple dendrite ablation. **(a)** Peak-normalized tuning curves of the neuron with three sequential basal dendrite ablation. The first ablation had little effect. The second ablation caused the cell’s tuning curve to shift by ~20 degrees. After the third ablation the tuning curve shift of the main peak remained unchanged. **(b)** Z projections depicting soma and basal dendrites showing structure of example neuron before ablation (left) after 1 basal ablation (middle) and after 2-basal+apical ablation (right). Red arrow points the basal dendrite that ablated first. Pink arrow points the secondly ablated basal dendrite. Apical dendrite ablation was assessed in other z-stack images (Movie S4) and apical dendrite is not depicted in these images. **(c)** peak-normalized tuning curves of the neuron after two basal dendrite followed by apical dendrite ablation.

**Supplementary Figure 10.**
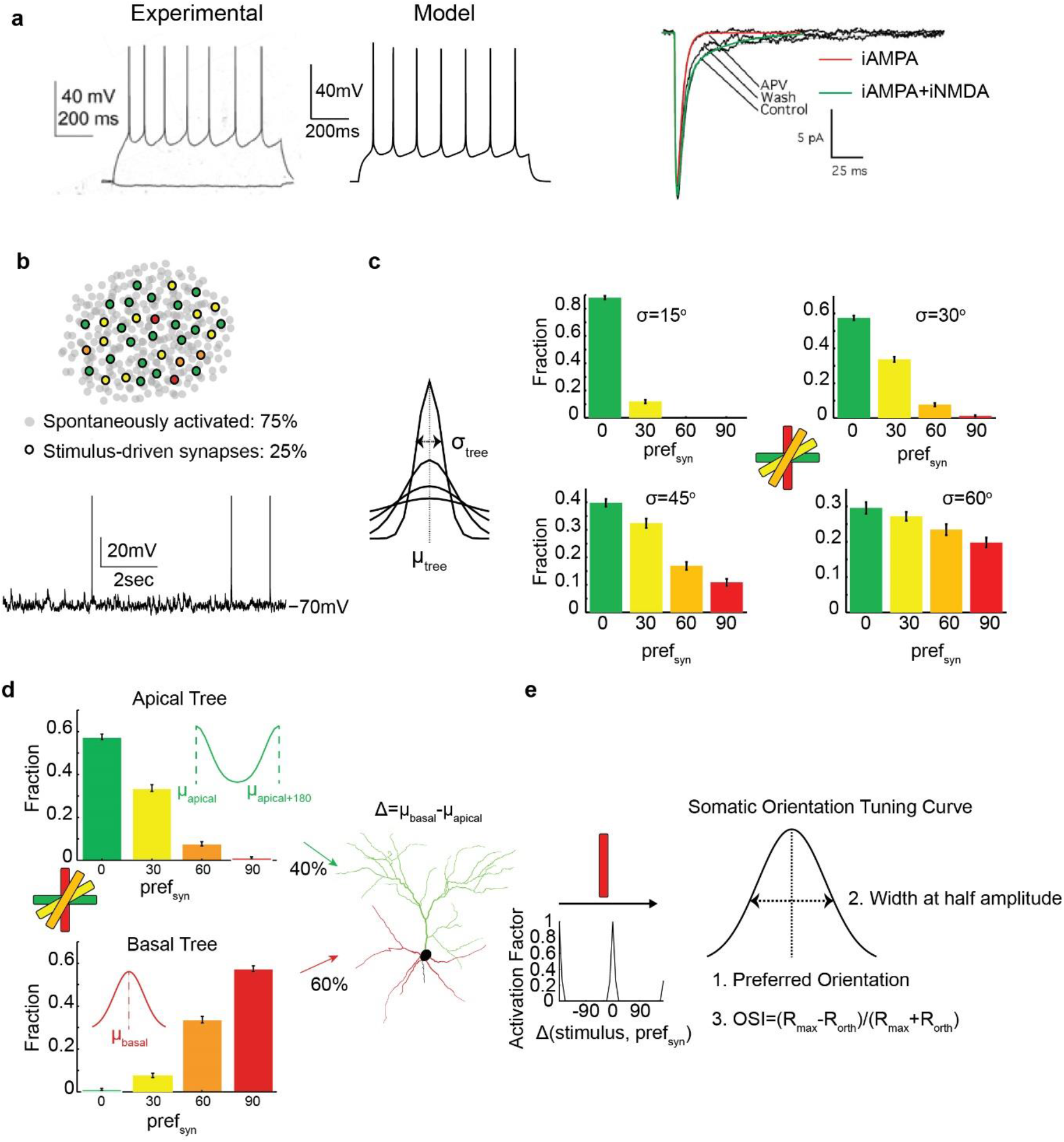
Single neuron model of orientation preference. **(a)** Experimental (left) ^43^ and model (right) responses to 900ms current clamp (0.16nA for both cases). Far right: Experimental ^41^ (black traces) and modelling properties of mEPSCs under control conditions (combined AMPA and NMDA current, green trace) and in the presence of APV (modelled as iAMPA, red trace). **(b)** *<u>Top</u>*: From the synaptic pool, 25% were stimulus driven (colored dots) and the rest were activated from background noise ^8^. *<u>Bottom</u>*: Indicative trace showing fluctuations of the membrane potential in the presence of background synaptic activity. The membrane potential rests above −70mV. Spikes are truncated for visualization purposes. **(c)** Each tree was characterized by a μ_tree_±σ_tree_ that determined the individual orientation preferences of activated synapses. Right: Fraction of synapses with a specific orientation preference (pref_syn_), for σ_tree_=15°, 30°, 45° and 60°. Bin width corresponding to the value reported on the x-axis is ±10°. **(d)** Exemplar case of the simulation setup, for **Δ(μ_basal_,μ_apical_)** =90°, showing the distributions of pref_syn_ of synapses for the apical (top) and basal (bottom) trees. Note that in this example, the dendritic trees have the same σ, but different μ. Synapses of each tree are assigned with a preferred orientation and then uniformly distributed along the apical and basal dendrites. **(e)** *<u>Left</u>*: When a stimulus is ‘presented’, the activation pattern of each synapse depends on the difference of pref_syn_ and the presented stimulus (activation factor). *<u>Right</u>*: Metrics of the resulting tuning curve at the soma.

**Supplementary Figure 11.**
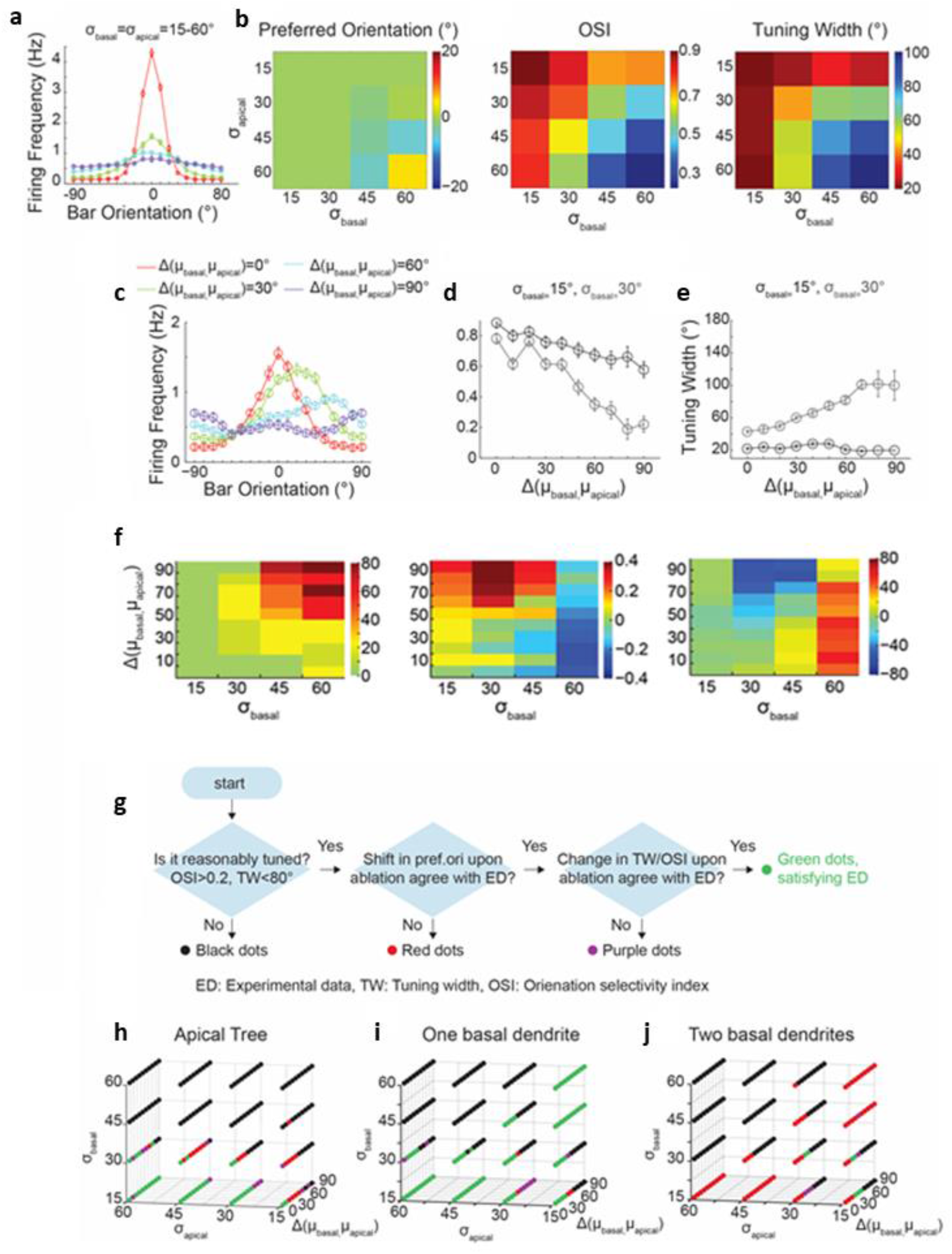
Input structures that replicate the experimental results. **(a)** Average orientation-tuning curves of 10 neurons for different σ. Δ(**μ_basal_,μ_apical_**) =0° across four conditions. σ_apical_=σ_basal_=15° (red), σ_apical_=σ_basal_=30° (green), 45° (blue) or 60 (purple). **(b)** Heat map plots of preferred orientation (left), OSI (center) and tuning-width (right) of the model neuron for different σ_basal_ and σ_apical_. Δ(**μ_basal_,μ_apical_**) was fixed at 0° and μtrees was arbitrarily set at 0°. **(c)** Average orientation-tuning curves of 10 neurons with varying Δ(**μ_basal_,μ_apical_**), (σ_apical_=σ_basal_=30°). **d-e**, OSI **(d)** and tuning-width **(e)** as a function of Δ at σ_basal_=15° (black) and σ_basal_=30° (grey). For both σ_apical_=30°. Error bars are SEM. Note that plots in **a-e** are from model neurons that are intact, not ablated. **(f)** Heat map depiction of pre- and post-apical dendrite ablation difference in preferred orientation (left), OSI (center) and tuning-width (right) when **σ_apical_**=30°, for **σ_basal_**={15°, 30°, 45°, 60°} and **Δ(μ_basai_,μ_apicai_)**={0°,…,90°}. **(g) Logic diagram of categorizing simulation results for 3D heat map figure h-j.** *<u>Black dots</u>:* parameter space that results in tuning curves with ÜSI<=0.2 and tuning-width >=80° before or after ablation in more than 30% of the simulated neurons. If differences in preferred orientation before and after ablation agree with experiments (i.e. <10° mode shift for apical and single basal cuts and ≥10° mode shift for two basal cuts), they are further categorized as green or purple dots based on their post ablation tuning-width and OSI. Thresholds to classify a change in OSI or tuning-width correspond to the mean+1std of the experimental data and are mean OSI change>0.2 and mean tuning-width change>10°. Model neurons that did not agree with experimental shifts in orientation preference are shown in red. **h-j** Summary of dendrite ablation model results for apical (h), single basal (i) and double basal (j) dendrite ablation. See (g) for logic of categorizing simulation results. For each simulated neuron (n=10), every combination of one or two basal dendrites was removed. **(i)** and **(j)** *<u>Green dots</u>:* mode shift in preferred orientation is *<u>Red dots</u>:* mode shift in preferred orientation is ≥10°. *<u>Purple dots</u>:* no mode shift change in preferred orientation WITH changes in tuning-width or OSI (see (g)). **(j)** *<u>Green dots</u>:* mode shift in preferred orientation is <10° AND tuning-width /OSI remain unchanged upon two basal dendrite cut. Max mode shift in preferred orientation among green dot conditions is 10°. *<u>Red dots</u>:* no shift in the preferred orientation (*<u>Purple dots</u>*: mode shift in the preferred orientation (≥10°) WITH changes in tuning-width or OSI (see legends for (g) for detail). In all h-j, only the green dots satisfy the experimental data. Note that there is no parameter space combination that simultaneously satisfies the experimental data of apical tree, one basal and two basal dendrites ablation. Therefore, this necessitates the introduction of differentially tuned basal dendrites. See Figure 5 and manuscript.

**Supplementary Figure 12.**
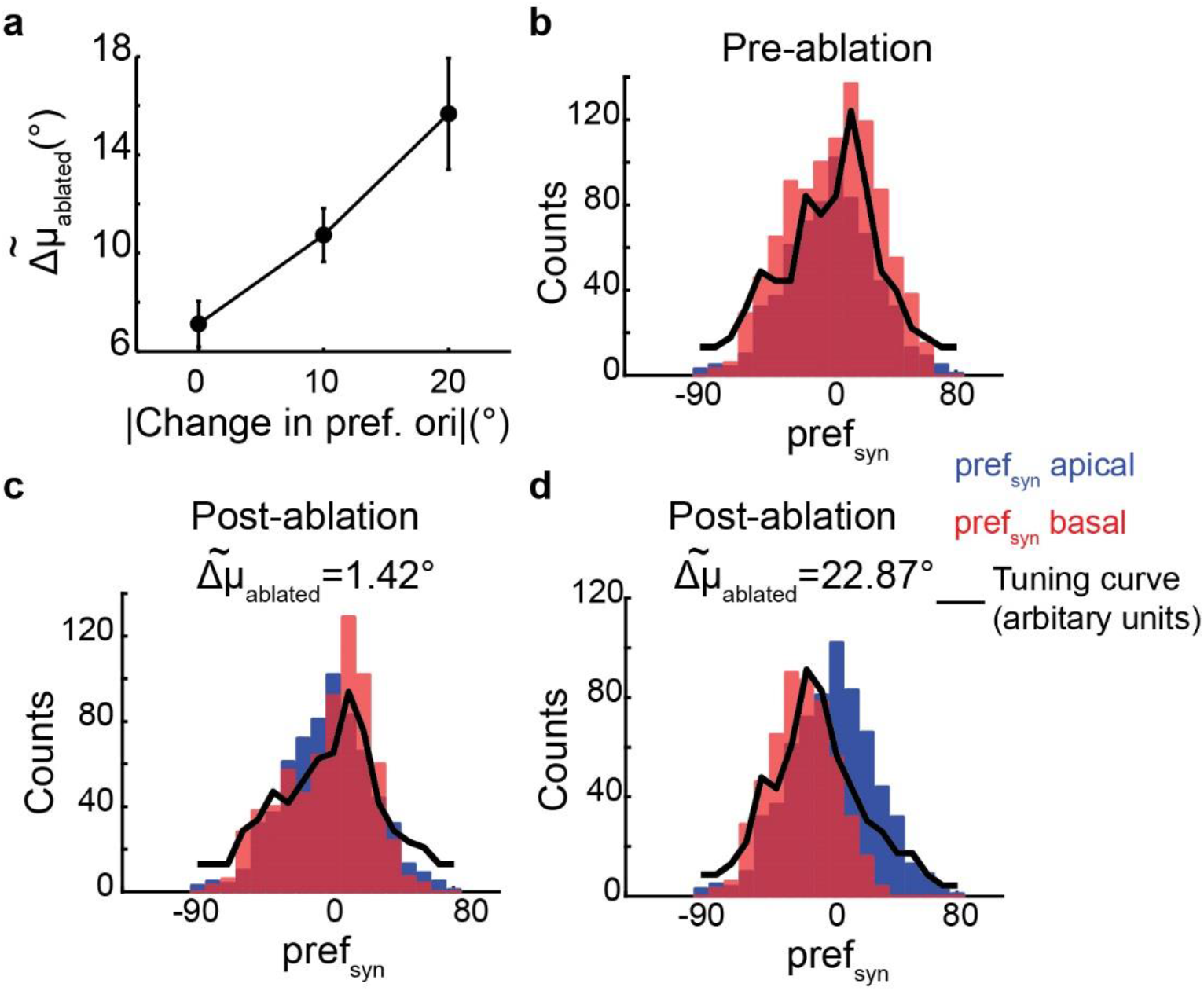
Mean orientation preference change in basal dendrites after two basal dendrite ablation correlates with preferred orientation change at the soma (in the model). **(a)** The change in length-normalized mean orientation preference of the input across basal dendrites following two basal dendrite ablation (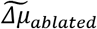; see Supplementary methods) as a function of the change in preferred orientation of the model neurons. Note that, as expected, they are highly correlated. Simulation parameters: Δ_μ_=40°, σ_apical_=30°, σ_basal_= 15°. Error bars: SEM. **(b)** Histogram of single synaptic preferred orientation (pref_syn_) of basal (orange) and apical dendritic trees (blue) of an exemplar neuron. Overlap appears as dark red. *<u>Black Curve</u>*: Resulting tuning curve of this neuron (arbitrary units). **(c)** For the same simulated neuron, removing two dendrites, resulting in small 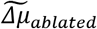 (1.42°) does not alter orientation-tuning. **(d)** However, removing two dendrites resulting in large 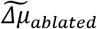 (22.87°) alters both input structure and orientation-tuning. 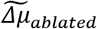 significantly increased with increasing post-ablation shift in orientation preference (p=0.0087, Kruskal-Wallis test)

**Supplementary Figure 13.**
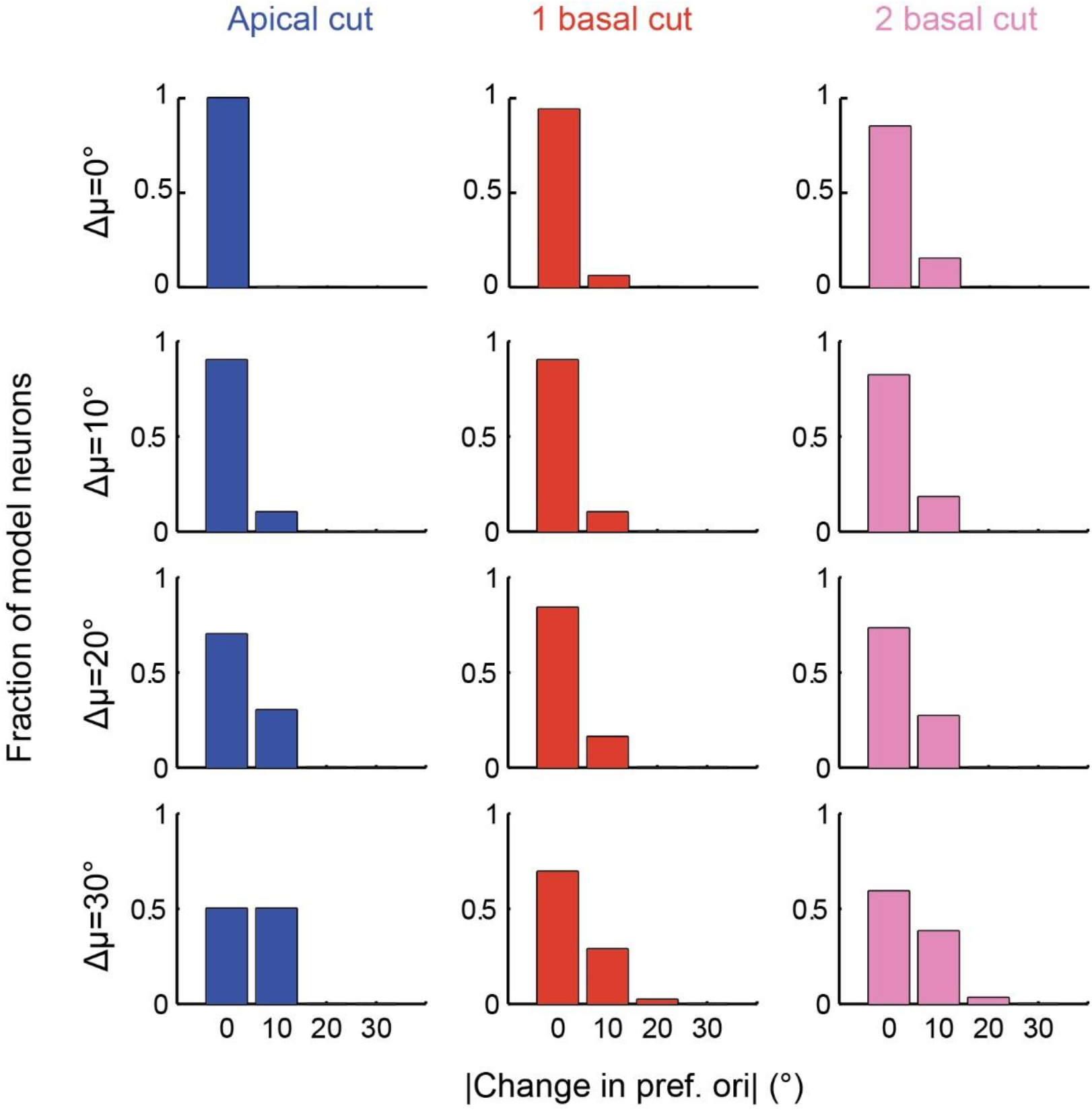
Biophysical model with broadly tuned basal dendrites (drift model). Figure in the format of Figure 5b but for σ_basal_=30°. Again, σ_apical_ is fixed at 30°. Gradually increasing Δμ leads to a very small shift in orientation preference with 2-basal cut that is a worse fit to experimental data than σ_basal_=15°. For all, mean change in tuning-width is < 10° and mean change in OSI is < 0.2 (thresholds correspond to the mean+1std of the experimental data). Disparities of 40°or greater fail to generate tuning curves comparable to experiment (0SI≤0.2, tuning-width ≥80° for more than 30% of simulated neurons). For each model neuron (n=10 per **Δμ**), all combinations of one or two basal dendrite cuts were simulated.

**Supplementary Table 2.** Active conductances of model neuron

| Conductance (mS/cm<sup>2</sup>) | Soma | Apical | Basal |
| --- | --- | --- | --- |
| $g_{Na}$ | 0.505 | 0.303 | 0.303 |
| $g_{Kdr}$ | 0.05 | $1.5 \cdot 10^{-3}$ | $1.5 \cdot 10^{-3}$ |
| $g_{Km}$ | $2.8 \cdot 10^{-3}$ | $1.27 \cdot 10^{-3}$ | $1.27 \cdot 10^{-3}$ |
| $g_A$ | 5.4 | diameter$\leq 0.8\mu m$: 108<br>diameter$\leq 0.8\mu m$: 10.8 | diameter$\leq 0.8\mu m$: 108<br>diameter$\leq 0.8\mu m$: 10.8 |
| $g_T$ | 0.03 | $x \leq 260\mu m$:<br>$0.029 \cdot \sin(0.009 \cdot x + 0.88)$<br>$x > 260\mu m$: 0.012 | $0.03 + 6 \cdot 10^{-5} \cdot x$ |
| $g_{HVA}$ | $0.05 \cdot 10^{-3}$ | $x \leq 260\mu m$:<br>$0.049 \cdot 10^{-3} \cdot \sin(0.009 \cdot x + 0.88)$<br>$x > 260\mu m$: $0.02 \cdot 10^{-3}$ | $0.05 \cdot 10^{-3} + 10^{-7} \cdot x$ |
| $g_{KCa}$ | $2.1 \cdot 10^{-3}$ | $2.1 \cdot 10^{-3}$ | $2.1 \cdot 10^{-3}$ |

**Supplementary Table 3.** Passive and active properties of the model neuron.

|  | Model | Cho et al., 2010 |
| --- | --- | --- |
| RMP, mV | -79 | -78.56±1.34 |
| IR, MΩ | 123.6 | 125.2±8.2 |
| $\tau$, ms | 17.3 | 16±0.7 |
| AP amplitude, mV | 66.1 | 67.8±1.8 |
| AP threshold, mV | -41.8 | -37.7±1.3 |
| AHP, mV | 17.9 | 13.3±0.5 |
| P-T time, ms | 38.6 | 55.3±2.7 |
| AP adaptation | 1.16 | 1.18±0.02 |

RMP: resting membrane potential, IR: Input Resistance measured at hyperpolarizing current (-0.04nA), AP: action potential, AHP: after hyperpolarization measured at depolarizing current (0.16nA), P-T peak-trough.

**Supplementary Table 4.** Synaptic parameters.

| | Conductance (nS) | $\tau_1$, ms | $\tau_2$, ms |
| --- | --- | --- | --- |
| NMDA | 1.15 | 2 | 30 |
| AMPA | 0.84 | 0.1 | 2.5 |
| GABA<sub>A</sub> | 1.25 | 0.2 | 1.4 |

**Supplementary Movie 1**. Animated Z-stacks (1 micron step size) of an example neuron going through an apical dendrite ablation. The movie starts from bottom-up and then reverses to go top-down to the same point before stopping. It is best that each panel is seen separately first, before comparing. *<u>Left Panel:</u>* The neuron structure prior to ablation. Arrow shows the point on the apical trunk targeted for ablation. *<u>Second Panel</u>:* Neuron has been subjected to a single point scan targeted to the location shown by the arrow. Note that the neuron and its dendrites increased in brightness (shown here 10 minutes following the 1^st^ point scan), likely as a result of calcium influx secondary to injury. However in this case the injury did not definitively sever the dendrite, which appears still grossly intact. Accordingly, the neuron regained its baseline fluorescence after ~30 minutes and the apical tuft dendrite remained intact (not shown here). *<u>Third Panel</u>:* The neuron is shown here 10 minutes following a second point scan targeted to the same location (shown by the arrow). Note that there is an interruption in fluorescence and the distal dendritic branches exhibit early beading morphology (a clear sign of successful ablation). Neurons whose dendrites were successfully ablated exhibited immediate but transient (<20 min) increase in fluorescence that included the targeted dendritic branches and the soma. The targeted dendritic branches then typically assumed a beads-on-a-string appearance prior to the disappearance of fluorescence. *<u>Right Panel</u>:* The neuron is shown 5 days following ablation. Note that the apical dendrite has now entirely disappeared. Note also that several other dendritic branches that were close (well within a 5μm range; see also Fig. 1c) but not connected to it, remain intact.

**Supplementary Movie 2**. 3D projection images of an example neuron pre-(left) and 5 days post-(right) apical dendrite ablation.

**Supplementary Movie 3**. Animated Z-stacks (moving coronally) of an example ablated neuron that was immunostained with anti-GFP (green) and anti-tuj 1 (magenta). More detailed information is described in Supplementary Figure 3. The arrow indicates the ablation point. Note that there is no discernible disorganization of the nearby anti-tuj 1 stained neuropil (the areas of absent staining appear to reflect nearby neurons that were not GFP stained). This is in agreement with a prior electron microscopy study (see Fig 2 of ^12^) using a similar protocol, which demonstrated that the laser ablation lesion is highly contained within an area ~5 microns in diameter.

**Supplementary Movie 4**. Animated Z-stacks (1 micron step size) of an example neuron that underwent 2 basal dendrite ablation followed by an apical dendrite ablation. The movie starts from bottom-up and then reverses to go top-down to the same point before stopping. It is best that each panel is seen separately first, before comparing. *<u>Left Panel</u>:* the neuron prior to ablation. Note the red and purple arrow pointing to the basal dendrites to be ablated, and the cyan arrow pointing to the apical trunk origin near the soma. *<u>Middle Panel</u>:* Picture of the same neuron after two basal dendrites have been ablated. The red and purple arrows point to the ablation targets, showing the targeted basal dendrites have been severed from the soma. Beaded remnants of their distal branches can be seen along their prior trajectories in the vicinity of the cell. Note that the cell is highly fluorescent as the Z-stack was obtained during the ablation of the apical dendrite. *<u>Right Panel</u>:* The same neuron 5 days following apical dendrite ablation. The cyan arrow indicates the apical dendrite ablation point. Note that the apical dendrite has deteriorated (the visible branches that can be seen towards the top a little below the arrow represent basal dendrites whose insertion point was close to that of the apical dendrite). Despite having lost 2 basal and the apical dendrites the cell survived and remained visually responsive and orientation-tuned (Supplementary Fig. 6c).

## Supplementary Data

### Model neurons

The morphologically detailed L2/3 V1 pyramidal neuron model (Figure 4A) was implemented in the NEURON simulation environment ^36^. Model neurons are available in ModelDB (private model until publication, https://senselab.med.yale.edu/modeldb/enterCode.cshtml?model=231185, password: nassi123). The passive properties of the model neuron were: membrane capacitance (C_m_) 1 μF cm^-2^, membrane resistance (Rm) 11000 Ω cm^2^ and axial resistance (Ra) 100 Ω cm. In the basal and apical dendrites, C_m_ was doubled to account for dendritic spines. The resting membrane potential was set at −79mV and the input resistance (R_IN_) at 124 MΩ ^19,37,38^. Membrane time constant was 17 ms ^38^.

The active properties of the model were adapted from^18,19^. The model included conductances for fast voltage-dependent sodium channels (g_Na_), delayed rectifier potassium channels (g_kdr_), slow voltage-dependent potassium channels (g_M_), A-type potassium channels (g_A_), calcium-activated potassium channels (g_KCa_) and high and low voltage-activated calcium channels (g_HVA_ and g_T_ respectively). In all compartments a calcium buffering mechanism was included (τ=50ms). At the apical dendrites the conductances of the calcium channels increased for distances <=80 μm and then decreased and reached minimal levels for distances above 260 μm. At the basal dendrites, calcium channel conductances increased linearly ^39^. Compared to the conductance of the thin dendrites (diameter<=0.8), the conductance of the A-type potassium channel was 5% at the soma and 10% at the thick dendrites ^40^. Supplementary Table 2 shows the ionic conductances used in each compartment. Supplementary Table 3 lists the electrophysiological characteristics of the model neuron in comparison to experimental data of ^38^. Supplementary Fig. 7a illustrates the electrophysiological profile of the model neuron in comparison to an experimental trace adapted from ^43^ in response to the same current step pulse (0.16 nA). The model is in great agreement with all of the abovementioned experimental data.

Synaptic mechanisms included AMPA, NMDA and GABA_A_ synapses. Kinetics of miniature AMPA current (activation of 1 synapse) were fitted to experimental data of^41^. In particular, the model neuron was voltage clamped at −70mV and consecutively 1 AMPA synapse was activated at each dendritic branch. The average response of the model (red trace) as well as the experimental average response (in the presence of APV to isolate the AMPA current) is shown in the right panel of Fig S7A. The kinetics of the NMDA current were also fitted to the experimental data of^41^ (Supplementary Fig. 7a, green trace) in the absence of Mg++ blockade, to simulate the experimental absence of Mg^++^ (only for validation purposes). The peak amplitude of the NMDA current was set so that the ratio of the iNMDA (measured at +50mV) to iAMPA current (measured at −50mV) was 1.2^41^. This resulted in gNMDA=1.37*gAMPA. Miniature EPSC amplitude (iAMPA and iNMDA) was set at 20 pA^41^. The unitary IPSP (activation of 15 inhibitory synapses) was 0.8 mV measured at resting membrane potential −65mV and the duration at half amplitude was set to 15ms^44^. Properties of synaptic currents used are shown in Supplementary Table 4.

To estimate the total number of excitatory synapses, we assumed synaptic density of 2 excitatory synapses (spines)/μm^15,45^. Excitatory synapses consisted of both AMPA and NMDA conductances. The total dendritic length of the model neuron is 3298μm and thus the total number of excitatory synapses was 6596, that is within the range of the experimentally reported total number of excitatory synapses in layer 2/3 visual cortex^15^. 60% of these were randomly distributed to the basal dendrites and 40% to the apical dendrites^15^. Distribution of synapses to individual dendritic compartments was adjusted so that each dendrite had the same synaptic density, that is, the number of synapses each dendrite received was proportional to its length. 15% of the total number of synapses were inhibitory^2,15^. 7% of inhibitory synapses were located at the soma, 60% at the basal dendrites and 33% at the apical dendrites 42. 25% of the total number of excitatory synapses were stimulus-driven^8^. The rest 75% of excitatory synapses, as well as the inhibitory synapses^46^, were independently driven by Poisson spike trains with mean frequency 0.11Hz (Supplementary Fig. 7b). This resulted in spontaneous spiking activity at frequency 0.28±0.37 Hz that is within the experimental range from in vivo recordings in mice^20^.

Each stimulus-driven synapse was tuned (i.e. had a preferred orientation, pref_syn_) in one of the thirty-six different angles (0°-350° with step 10°). Tuning was not direction selective i.e. a synapse with tuning at 0° was also tuned at 180°. The orientation-preference distribution of the stimulus-synapses followed the sum of two Gaussians centered on the mean orientation preference (orientation preference with maximum probability) of the dendritic tree (μ_tree_). We varied the standard deviations (σ_basal_, σ_apical_) of the distributions from 15° to 30°, 45° and 60° (Supplementary Fig. 7c). For σ=30°, distributions resemble the ones reported in^8^. In addition, we ranged the difference (Δ) of μ_basal_ and μ_apical_ from 0° to 90°, with step 10° (by varying the μ_basal_ and arbitrary keeping constant the μ_apical_ at 0°, Supplementary Fig. 7d). In the drift model, synapses of each basal dendrite (and not the whole dendritic tree) were assigned a preference according to the respective μ_b_, σ_b_ (Fig 5A). The μ_b_ of each dendrite was chosen from a predefined range (Δ_μ_).

60% of orientation-tuned synapses were randomly dispersed to basal dendrites and the rest 40% to the apical dendrites^4,15^ (Supplementary Fig. 7d), so that all dendrites have the same density of stimulus synapses. Stimulus (a bar at a specific direction) was ‘presented’ after an initial period of 500ms and lasted for 2 secs as in the experimental setup (Fig 4b). During the ‘presentation’ of the moving bar (0°-180°, with step 10°), stimulus-driven synapses were activated by Poisson spike trains (Supplementary Fig. 7e). Each stimulus-driven synapse received a spike train with mean 0.3 Hz*activation factor, where the activation factor depended on the Δ(stimulus, pref_syn_) (Supplementary Fig. 7e). The neuron responded under the control conditions (see below) for the preferred orientation with 1.65 ±0.94 Hz^20^. Control condition was the one corresponding to σ_apical_=σ_basal_=30°, μ_apical_=μ_basal_=0°, Δμ=0°. Under this condition, the OSI was 0.78 and the tuning width 43° (Fig. 4c), replicating the experimental data for orientation tuning in L2/3 neurons of the visual cortex.

Simulation of the apical tree and basal dendrite ablation was performed by removing the respective compartments. We simulated basal dendrite cutting by removing each one of the five primary basal dendrites for one dendrite cutting or all combinations of two primary basal dendrites for two dendrites cutting. The model neuron’s firing rate increased following apical dendrite ablation due to a 2.5-fold increase in input resistance (as seen in^47^), while basal dendrite ablation slightly decreased firing rate due to excitatory input loss. In the model, synaptic scaling (reduction) of excitatory transmission was implemented (as observed in prior studies of homeostatic plasticity^48^) to normalize the neuron’s firing rate without affecting the tuning curve shape. Specifically, pre-/post-ablation excitatory synaptic weights (gAMP and gNMDA) were adjusted to result in the same levels of spiking activity at the preferred and the orthogonal orientation under control condition, as indicated by the calcium data (Supplementary Fig. 4).

For each condition, we simulated 10 model neurons (different in respect to the pref_syn_ of each synapse, as well as in their location along the trees) and for each neuron we averaged the resulting spiking activity over 10 repetitions. For each condition, preferred orientation at the cell body was defined as the orientation (0° to 170° with step 10°) to which the neuron displayed the highest mean firing frequency (Supplementary Fig. 7e). Since we did not model direction selectivity, each preferred orientation was also represented by its symmetric angle (e.g. 0° and 180°). Accordingly, the orientation tuning-width was the width at half amplitude of the fitted curve. Selectivity of orientation tuning for each condition was assessed using the OSI metric, defined as 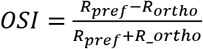. Simulated neurons that lacked realistic tuning curves (OSI ≤ 0.2, tuning width ≥ 80°) were excluded from the analysis. We further did not consider orientation tuning for combinations of simulation parameters in which more than 30% of the simulated neurons were categorized as not tuned.

In Figure 4F, mode change was plotted for orientation preference and average change was used for plotting changes in tuning-width and OSI.

In the drift model, Δμ of 50° or greater failed to generate tuning curves comparable to experiment (>30% of simulated cells), so the analysis was limited to Δμ=20-40°.

To assess if the change in input structure across basal dendrites following two basal dendrite ablation in fact relates to the change in orientation preference of the model neuron, we calculated 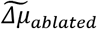, the difference between length normalized orientation preference in input structure across basal dendrites before and after two basal dendrite ablation. The 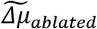 was calculated for each combination of ablated dendrites as follows:

For each primary basal dendrite,

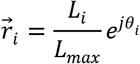

where L_i_ is the length of the i_th_ dendrite normalized to the length of the longest dendrite (L_max_) and θ_i_ the mean of orientation preference of the distribution of synapses of the respective basal dendrite (in radians). The population vector, using Euler’s formula, is:

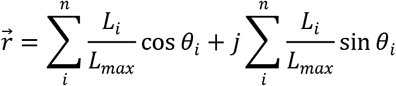

where n is the number of primary basal dendrites. The angle of the population vector is:

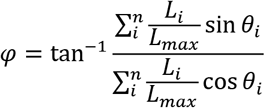

Finally, we defined 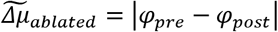, where ϕ_pre_ is the population vector of the input structure before the ablation of the primary basal dendrites and ϕ_post_ after (in degree). Simulations were performed in the Computational Biology Lab High Performance Computational (HPC) cluster consisting of 312 High Performance CPU cores and 1.150 Gigabytes of RAM. Analysis was performed in Python 2.7.13 via Anaconda 4.4.0. The model neuron code is available in ModelDB (accession number 231185).

